# Non canonical scaffold-type ligase complex mediates protein UFMylation

**DOI:** 10.1101/2022.01.31.478489

**Authors:** Joshua J. Peter, Helge M. Magnussen, Paul Anthony DaRosa, David Millrine, Stephen P Matthews, Frederic Lamoliatte, Ramasubramanian Sundaramoorthy, Ron R Kopito, Yogesh Kulathu

## Abstract

Protein UFMylation is emerging as a posttranslational modification essential for endoplasmic reticulum and cellular homeostasis. Despite its biological importance, we have a poor understanding of how UFM1 is conjugated onto substrates. Here, we use a rebuilding approach to define the minimal requirements of protein UFMylation. We find that the reported E3 ligase UFL1 is inactive on its own and identify UFBP1 to bind UFL1 to form an active E3 ligase complex. While UFC1 is an intrinsically Cys-reactive E2, we do not identify any catalytic cysteines on UFL1/UFBP1, suggesting a scaffold-type E3 ligase mechanism. Interestingly, the E3 ligase complex consists of winged-helix (WH) domain repeats that activate UFC1 for aminolysis. We identify the adaptor protein CDK5RAP3 to bind to and regulate E3 ligase activity potentially by preventing off-target UFMylation. In summary, our work identifies the minimal requirements for UFMylation and reveals regulatory principles of this atypical E3 ligase complex.

## Main

UFM1 is a ubiquitin-like (Ubl) modifier that contains the characteristic β-grasp fold found in all UBLs and is highly conserved in multicellular eukaryotes ^1,2^. Like other UBLs, UFM1 is covalently attached to lysine residues on substrates via an enzymatic cascade involving an E1 activating enzyme, UFM1-activating enzyme 5 (UBA5), E2 conjugating enzyme, UFM1-conjugating enzyme 1 (UFC1), and an E3 ligase, UFM1 E3 ligase 1 (UFL1) ^3,4^. Recent studies have identified the ribosomal protein RPL26 to be the principal target of UFMylation which plays a critical role in endoplasmic reticulum (ER) homeostasis ^5–7^. Besides ER associated roles, UFMylation has also been implicated in other cellular processes including protein translation, DNA damage response and nuclear receptor mediated transcription ^8–13^. Further, loss of or mutation of components of the UFMylation machinery has been linked to many diseases such as cancer, type-2 diabetes, neurological disorders and cerebellar ataxia, where failure of ER homeostasis and protein quality control could be one of the major contributing factors ^12,14–17^. Indeed, UFMylation is an essential posttranslational modification for animal development as loss of UFM1 or any of the UFMylation enzymes results in failure of erythropoiesis and embryonic lethality ^18–21^.

UFM1 is expressed in a precursor form consisting of 85 amino acids. It is then post-translationally processed by UFM1-specific proteases (UFSPs) which cleave off the last two residues at the C-terminus to generate mature UFM1 that contains an exposed C-terminal glycine, Gly^83 22^. Mature UFM1 is activated by the E1, UBA5 via the formation of a high-energy thioester bond between Cys^250^ of UBA5 and Gly^83^ of UFM1 ^3,23^. Activated UFM1 is then transferred from UBA5 on to the catalytic Cys^116^ of UFC1 ^3,24^. UFC1 has a canonical UBC fold and in addition an N-terminal helical extension whose function is not known ^25,26^. UFC1 lacks many of the features conserved in many E2s such as the catalytic HPN motif which suggests unique modes of action and regulation. Further, ubiquitin (Ub) E2 enzymes such as UBE2D3 associate with Ub via a backside interaction which mediates E2 dimerization and processivity of polyUb formation ^27,28^. For UFC1, it is unknown if similar mechanisms exist to regulate its activity as detailed biochemical characterization is lacking.

UFL1 is the only E3 ligase identified to date in the UFMylation pathway catalysing the final step of transfer of UFM1 from UFC1 to substrates ^29^. Indeed, loss of UFL1 results in loss of UFMylation and mice lacking Ufl1 phenocopy loss of UFM1 exhibiting failure in haematopoiesis and embryonic lethality suggesting that it could be the main E3 ligase ^30^. While data implying UFL1 is the E3 ligase stem from knockout studies and overexpression studies, direct biochemical evidence demonstrating that UFL1 is an E3 ligase is insufficient ^24^. Consequently, the mechanism of how UFL1 functions as an E3 ligase is also not understood. In general, E3 ligases are classified either as scaffold-type ligases that bring together a UBL-charged E2 with substrate for direct transfer of the UBL from the E2 on to substrate, or as Cys-dependent enzymes where the UBL is first transferred from the E2 to the catalytic Cys of the E3 for subsequent transfer to the substrate ^31^. Examples of scaffold-type E3s are RING family ligases and RANBP2 whereas HECT, RBR and RCR enzymes typify the latter type ^32–36^. Intriguingly, UFL1 does not possess any conserved sequence or domain features found in known E3 ligases. Hence it is not clear if UFL1 is a Cys-dependent E3 enzyme or uses a scaffolding mechanism.

UFL1 is predominantly found anchored at the ER through its interaction with an adaptor protein, DDRGK1/UFBP1 (UFM1 binding protein-1) which has also been suggested to be one of the main substrates of UFL1 ^29,37^. UFBP1 localizes to the ER membrane via an N-terminal transmembrane segment and its function is unclear ^5^. Intriguingly, loss of UFBP1 affects the stability and expression levels of UFL1 ^20,29,37^. CDK5RAP3 is another poorly characterized protein commonly associated with UFL1. Several anecdotal reports have led to speculations that UFL1, UFBP1 and CDK5RAP3 together form an E3 ligase complex ^24,38^. Only a handful of substrates of UFL1 have been identified to date and many of them are ER localized ^38,39^. In addition to monoUFMylation, polyUFM1 chains can be assembled which are mainly linked via K69 ^12^. In the Ub system, the type of polyUb linkage formed is generally dictated by the last enzyme in the cascade that forms a thioester linkage with Ub before transfer to substrate. Scaffold-type E3s such as RING E3 ligases bind to charged E2 (E2∼Ub) to mediate transfer of Ub onto substrate, and hence the E2 enzymes are thought to determine the linkage type assembled. In contrast, catalytic Cys containing E3 ligases form an E3∼Ub thioester intermediate and dictate the linkage type formed ^31,34^.

Given the lack of understanding of the molecular mechanism underpinning the UFM1 conjugation machinery and the mechanisms governing its regulation, here we adopt a rebuilding approach with purified components to fully reconstitute UFMylation *in vitro*. Using this reductionist approach, we reveal that UFL1 on its own is an unstable protein that cannot support UFMylation. We demonstrate that UFL1 together with UFBP1 forms a functional hetrodimeric E3 ligase complex and provide in depth biochemical characterization of the ligase complex. Structure prediction using Alphafold provides an explanation for the underlying reason for complex formation between UFL1 and UFBP1 and offers a rational framework for making truncations which help establish the minimal regions on UFL1 and UFBP1 essential for ligase activity. Parallel investigations of UFC1 reveal that it is an intrinsically Cys reactive enzyme that is activated by the UFL1/UFBP1 ligase complex for aminolysis. Our analyses reveal UFL1/UFBP1 to be a scaffold-type E3 ligase. Further, we identify CDK5RAP3 to bind to the E3 ligase complex and inhibit UFMylation by preventing discharge of UFM1 from UFC1. Importantly, by reconstituting UFMylation of ribosomes *in vitro,* we find that CDK5RAP3 has a regulatory function as it restricts UFMylation towards *bona fide* substrates and lysine selectivity. Lastly, we identify an N-terminal helical extension on UFC1 to have an inhibitory role that is required by CDK5RAP3 to restrain UFMylation. Our mechanistic insights describe an atypical E3 ligase complex and provide a foundation for investigating the catalytic mechanism, regulation, and substrate specificity of this E3 ligase complex.

## Results

### UFC1 has intrinsic cysteine reactivity

The interaction of UFC1 with UBA5 and how UFM1 is transferred from UBA5 to the catalytic Cys is reasonably well understood ^23,24,40–42^. We therefore focussed on subsequent events and first analysed the intrinsic reactivity and stability of UFC1∼UFM1 (∼ denotes the thioester bond between UFM1^G83^ and UFC1^C116^) in comparison to the well characterized ubiquitin E2s, UBE2D3 and UBE2L3 ^43^. Intrinsic reactivity of E2s assayed using free amino acids can give insights into the ability of E2s to transfer UBLs onto substrates and can provide clues about the type of E3 ligase they work with and even the nature of the substrate ^27,36^. For instance, E2s such as UBE2D3 which function together with RING/scaffold like E3 ligases are capable of aminolysis and are therefore reactive towards the free amino acid Lys (**Fig 1A, B**) ^43^. On the other hand, UBE2L3 which functions only with Cys-dependent E3 ligases is incapable of aminolysis and lacks reactivity to free Lys on its own (**Fig 1B**) ^43^. When discharge of UFM1 from UFC1∼UFM1 onto free amino acids was analysed, we observed preferential discharge onto Cys while no discharge was observed in the presence of Lys, Arg, Ser and Thr (**Fig 1C**). Under the same experimental conditions, UBE2D3 discharged Ub in the presence of both Cys and Lys (**Fig 1C, Fig S1A**). Further, using increasing concentrations of free Lys or with prolonged incubation in a time course, we find that UFC1 has negligible Lys reactivity which is similar to UBE2L3 (**Fig 1D-E, S1B**). These results suggest that UFC1 has intrinsic Cys reactivity and may work with Cys-dependent E3 ligases.

**Figure 1:**
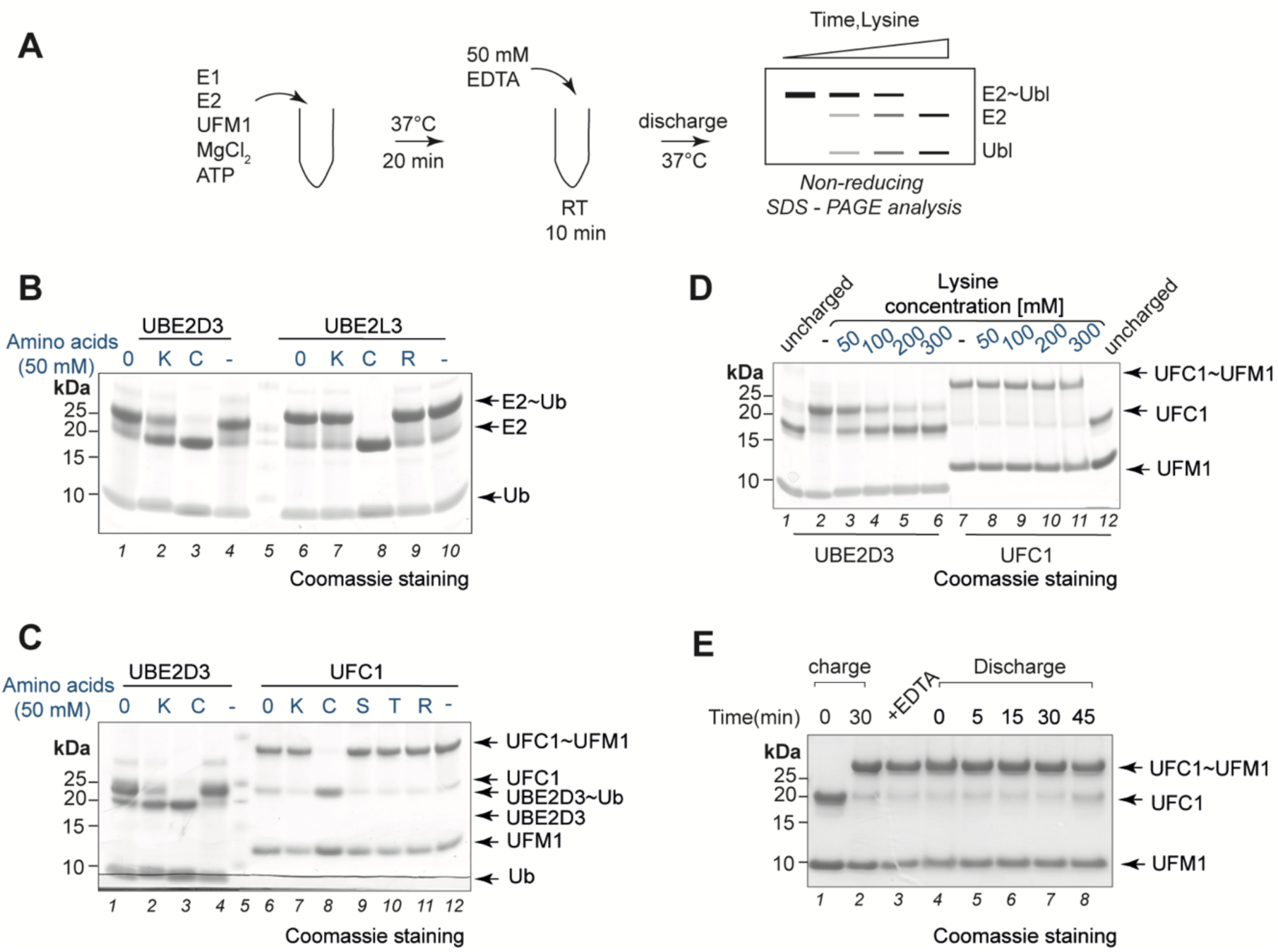
Nucleophilic reactivity profiling of UFC1. **A.** Schematic showing the workflow for substrate independent single turnover lysine discharge assays. **B.** Assay to monitor discharge of Ubiquitin from UBE2D3 and UBE2L3 in the presence of free amino acids. 0 indicates time 0. **C.** Assay to check for discharge of UFM1 from UFC1 in the presence of free amino acids. UBE2D3 is used as a positive control for lysine and cysteine discharge. **D.** Lysine discharge assays using UBE2D3 and UFC1 in the presence of increasing concentration of free Lysine. **E.** Time-dependent analysis of discharge of UFM1 from UFC1 in the presence of high concentration of free lysine (150 mM).

### UFL1 together with UFBP1 forms an active E3 ligase complex

To analyse how UFC1 works together with the E3 ligase UFL1 to transfer UFM1, we aimed to reconstitute UFMylation using UFL1. We expressed 6xHis-UFL1 in *E. coli* and purified it by affinity chromatography as the first step. Subsequent size exclusion chromatography (SEC) analysis indicated UFL1 to elute in the void fraction suggesting formation of soluble aggregates (**Fig S2A**). Our attempts to prevent aggregation by incorporating different solubility enhancing tags at the N-terminal or C-terminal ends of UFL1 or with different buffers and additives were unsuccessful. We therefore reasoned that UFL1 might require additional binding partners for its stability and activity. Indeed, several studies have shown that expressing subunits of protein complexes individually may result in aggregation ^44–46^. To identify binding partners of UFL1, we performed Yeast Two Hybrid (Y2H) screening using a cDNA library derived from human placenta as prey (**Fig S2B**). Surprisingly, only four hits were identified with UFBP1 being the most significant interactor. We therefore wondered if co-expressing UFBP1 with UFL1 might confer solubility and stability to UFL1. To achieve this, we used a co-expression system to express full-length UFL1 together with UFBP1 lacking its transmembrane sequence and purified the ligase complex with three enrichment steps (**Fig S2C**). Indeed, UFL1 co-expressed with UFBP1 lacking its N-terminal transmembrane segment (UFL1/UFBP1) is soluble, well behaved and no longer formed soluble aggregates in SEC (**Fig 2A**). We confirmed the formation of a stable UFL1/UFBP1 complex by analysing its precise molecular mass using a mass photometer ^47^ and SEC-MALS (**Fig 2B, S2D**). This revealed UFL1/UFBP1 to have a mass of 126 kDa which corresponds to that of a heterodimeric complex. Of note, the mass photometry measurements are performed at low nanomolar concentrations highlighting the stable nature of the complex.

**Figure 2:**
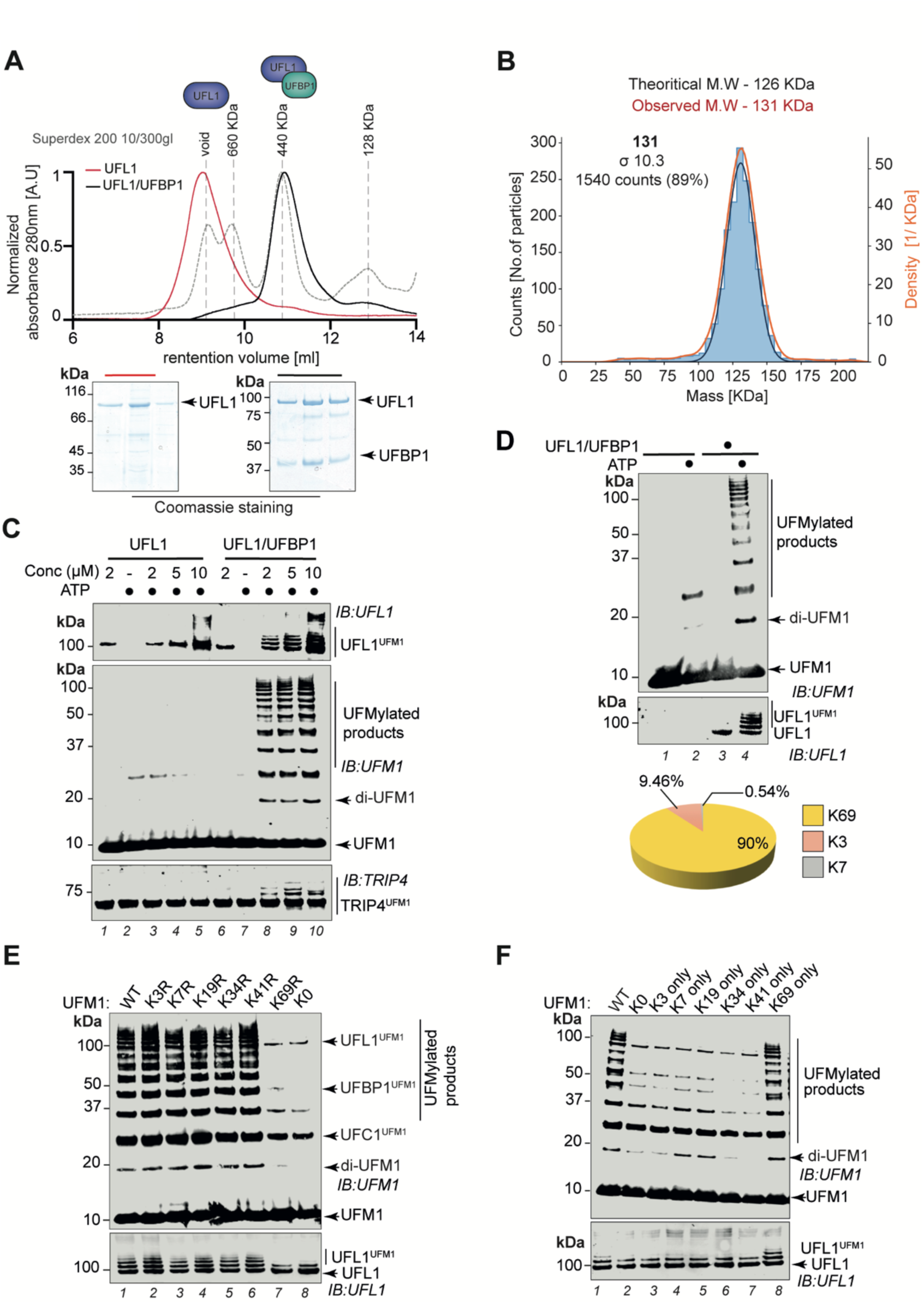
*In vitro* reconstitution of the active UFM1 E3 ligase. **A.** Comparison of size exclusion chromatography profiles of UFL1 expressed alone (Red) and co-expressed UFL1/UFBP1 complex (Black) run under identical buffer conditions on a HiLoad^TM^ Superdex 10/300 gl column. Molecular weight standards are shown in grey. Fractions corresponding to each peak were collected and analysed on 4-12% denaturing SDS-PAGE which is shown below. **B.** Mass photometry analysis of co-purified UFL1/UFBP1 complex. The theoretical and experimental molecular weights are indicated above. **C.** Immunoblot comparing UFL1 autoUFMylation *(Top)* and UFM1 chain synthesis *(Middle)* in the presence of UFL1 expressed alone and in complex with UFBP1. *(Bottom)* Assays to monitor UFMylation of substrates using purified TRIP4 in the presence of UFL1 alone and UFL1/UFBP1 complex. Reaction products were run on a 4-12% SDS-PAGE gel and analysed by immunoblotting using indicated antibodies. **D.** Immunoblot showing formation of free UFM1 chains in the presence and absence of UFL1/UFBP1. *(Bottom)* Schematic representation of the linkage composition of di-UFM1 chains obtained from LC-MS/MS analysis. **E.** *In vitro* UFMylation assay in the presence of Lys to Arg (K-R) and lysine less(K0) mutants of UFM1 to check for polyUFMylation *(top)* and UFL1 autoUFMylation *(bottom)*. **F.** *In vitro* UFMylation assay to check for formation of di-UFM1 chains in the presence of single Lys and Lys less mutants of UFM1.

Having obtained soluble UFL1/UFBP1 complexes, we next tested the catalytic activity of UFL1 expressed alone and in complex with UFBP1 using *in vitro* UFMylation assays containing UBA5, UFC1 and UFM1. Whereas UFL1 on its own did not show any appreciable activity, the UFL1/UFBP1 complex is an active ligase as evidenced by the formation of multiple UFMylated products and UFL1 autoUFMylation (**Fig 2C**). To check if UFL1/UFBP1 was capable of UFMylating substrates *in vitro*, we used TRIP4/ASC1, a protein reported to be UFMylated in cells ^12^. Indeed, UFL1/UFBP1 can efficiently UFMylate TRIP4 *in vitro* (**Fig 2C**). Intriguingly, exogenous addition of purified UFBP1 to UFL1 expressed on its own does not change the elution profile of UFL1 to match that of co-expressed UFL1/UFBP1 or restore its E3 ligase activity (**Fig S2E-F**). Taken together, these results show that UFL1 and UFBP1 form an obligate heterodimer required for E3 ligase activity.

Since polyUFMylated products are formed in UFMylation reactions, we next analysed the reaction products by mass spectrometry to determine the linkage types assembled. Interestingly, LC/MS analyses revealed that the polyUFM1 chains are formed which are predominantly linked via K69 with some K3 and K7 linkages also observed (**Fig 2D**). To validate these observations, we generated a series of K-R mutants where every Lys in UFM1 was individually mutated to Arg. While mutation of K3, K7, K19, K34 or K41 to Arg did not affect overall UFMylation, UFM1 K69R resulted in near complete loss of polyUFMylated species and unanchored diUFM1, closely resembling reactions with UFM1 K0 in which all Lys were mutated to Arg (**Fig 2E** lanes 7&8**, S2G**). Interestingly, western blotting for UFL1 revealed only monoUFMylated UFL1 in the presence of UFM1 K69R suggesting that UFL1 autoUFMylation involves formation of polyUFM1 chains on UFL1 that are linked via K69 (**Fig 2E**). We next generated K_only_ UFM1 mutants that contain only a single Lys while all others are mutated to Arg. UFMylation assays using these mutants confirm that K69 is the preferred linkage type assembled (**Fig 2F**). Single Lys-retaining mutants all abrogate polyUFMylation formation comparably to UFM1 K0 with the sole exception of K69-only UFM1 (**Fig 2F**). In reactions with UFM1 K69R or K0, mainly monoUFMylated products corresponding to modified UFL1, UFBP1 and UFC1 are formed. In summary, our reconstitution experiments reveal that UFL1 together with UFBP1 forms an active E3 ligase complex that can efficiently UFMylate substrates and assemble polyUFM1 chains that are K69-linked.

### UFL1/UFBP1 is a scaffold type E3 ligase complex activating UFC1 for aminolysis

Our analyses find that UFC1 is an intrinsically Cys-reactive E2 suggesting that UFL1 may be a Cys-dependent enzyme (**Fig 1**). The N-terminal region of UFL1 contains four Cys residues and sequence analysis show that all four vary in their degree of conservation with C32 being the most conserved (**Fig 3A**). Based on analysis of predicted folding propensity (**Fig S3A**) and secondary structure prediction, we made a C-terminal truncation of UFL1, UFL1^1–410^, which when expressed on its own formed soluble aggregates like full length UFL1 and required co-expression of UFBP1 to obtain a heterodimeric complex (**Fig S3B**). Importantly, this UFL1^1-410^/UFBP1 complex is an active ligase **(Fig S3C**). To identify the catalytic Cys in UFL1, we mutated each Cys individually to Ala in the UFL1^1-410^/UFBP1 complex. To our surprise, *in vitro* UFMylation assays showed that none of the single mutants had any impact on the autoUFMylation activity of UFL1 or UFMylation of substrates implying that UFL1 does not contain a catalytic Cys (**Fig 3B**). Since UFBP1 does not have any Cys residues, these results suggest that the UFL1/UFBP1 ligase complex may instead employ a scaffolding mechanism to transfer UFM1 on to substrate. Further, we do not observe any transthiolation products of UFL1/UFBP1 **(Fig S3D)**.

**Figure 3:**
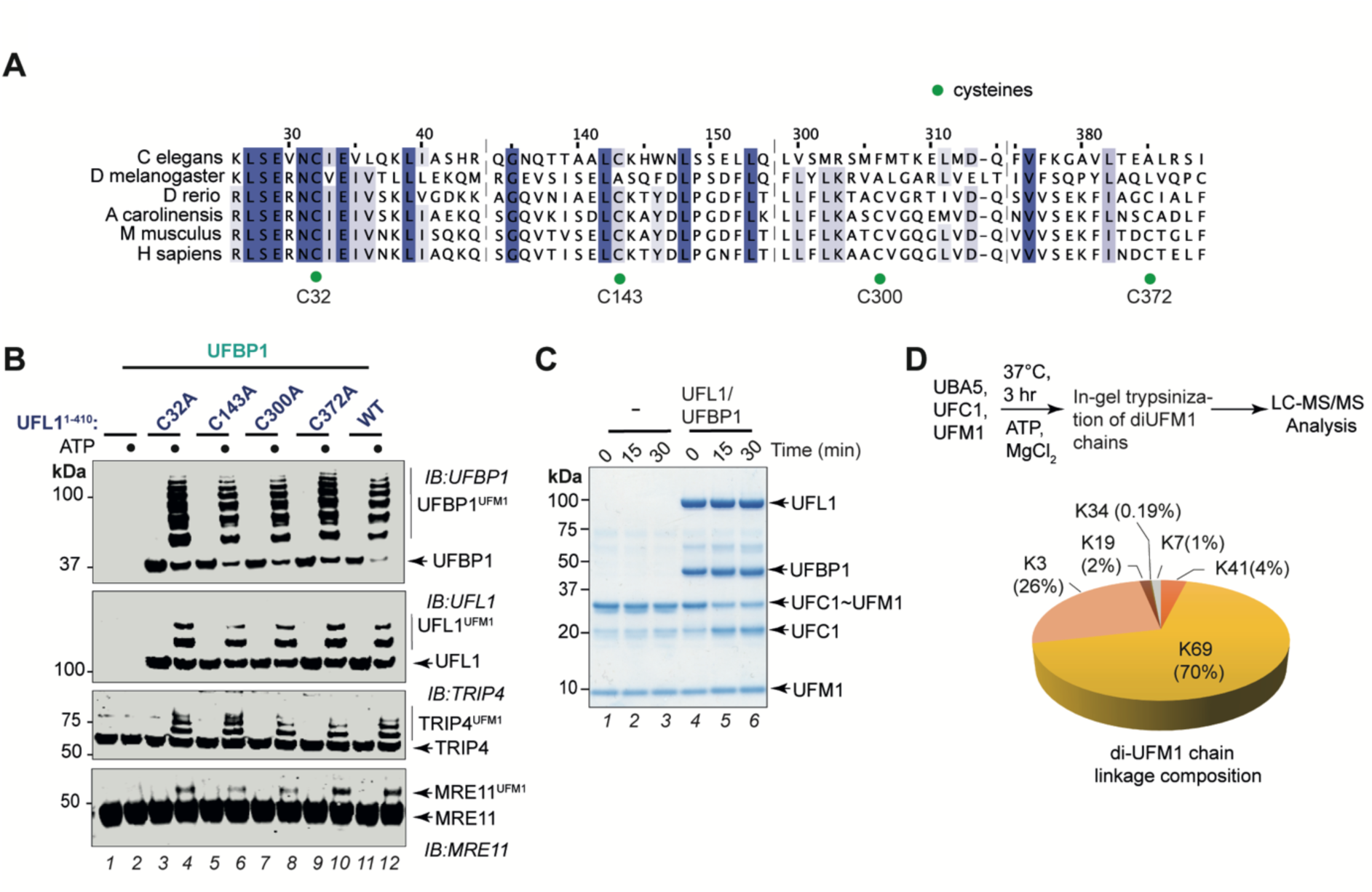
UFL1/UFBP1 is a scaffold type E3 ligase. **A.** Multiple sequence alignment of 1-410 a.a. region of UFL1 from various organisms to highlight the degree of conservation of cysteine residues in this region. **B.** Immunoblot showing *in vitro* UFMylation assays in the presence of C to A mutants of UFL1 to check for UFBP1 UFMylation *(top)*, UFL1 autoUFMylation *(middle)* and substrate UFMylation *(bottom)*. **C.** Coomassie stained SDS-PAGE gel showing discharge assays in the presence and absence of full length UFL1/UFBP1 complex. Charged UFC1 was incubated with lysine (50 mM) in the presence and absence of full length UFL1/UFBP1 complex and analysed for discharge on a 4-12% SDS PAGE gel under non-reducing conditions. **D.** Graphical representation showing the composition of linkage forms of di-UFM1 chains formed by minimal reconstitution of UBA5 and UFC1.

This raises the possibility that the UFL1/UFBP1 ligase complex could induce aminolysis of UFC1∼UFM1. Hence, we compared discharge of UFM1 from UFC1∼UFM1 onto Lys in the absence and presence of the ligase complex. Whereas UFC1 does not discharge UFM1 onto Lys on its own, UFM1 was readily discharged onto Lys in the presence of UFL1/UFBP1 (**Fig 3D**). Based on these observations, we suggest that UFL1/UFBP1 functions as a scaffold-type E3 ligase that binds to charged UFC1 to promote aminolysis.

While inefficient, UFC1 on its own can assemble free UFM1 chains which is significantly enhanced in the presence of the UFL1/UFBP1 ligase complex (**Fig S3E-F**). Interestingly, UFC1 assembles mainly K69 linkages (**Fig 3D**) and this linkage specificity is maintained in the presence of UFL1/UFBP1 (**Fig 2D**). Of note, this reveals that the linkage specificity is determined by the E2 and is not altered by the E3 ligase complex, a feature commonly observed in ubiquitin RING E3 ligases ^34,48,49^. These results further strengthen our conclusion that UFL1/UFBP1 is a scaffold-type ligase complex.

### Tandem WH domains of UFL1/UFBP1 constitute minimal E3 ligase

As UFL1 does not have any obvious sequence or domain features found in any of the known E3 ligases, we attempted to define the minimal catalytic region on UFL1. As our efforts to determine the structure of this complex were not successful, we used the recently released AlphaFold pipeline to predict the structure (**Fig 4A**) ^50^. Structure prediction of the complex had good predicted aligned error (PAE) scores (**Fig S4A**) and shows UFL1 and UFBP1 form a heterodimer (**Fig 4A-B**). Interestingly, UFL1 is predicted to contain 5 winged-helix (WH) domain repeats and a C-terminal helical domain (**Fig 4A** blue, **S4B-C**). The WH repeats are arranged such that a helix from each WH domain forms a helical backbone. The fifth WH domain on UFL1 is followed by a disordered region that extends into a stack of *α*-helices at its C-terminus which we refer to as CTR (C-Terminal Region). These WH domains could be classified as PCI-like WH domains, named so because it is commonly found in components of the proteasome lid, the COP9 signalosome and the eukaryotic translation initiator, eIF3 ^51,52^. In addition, UFL1 is predicted to have an N-terminal helix (1-25) followed by a partial or hemi-WH domain (pWH) (**Fig 4B**, blue). Following the transmembrane segment at its N-terminus, UFBP1 has a long coiled-coil region which we refer to as NTR (N-Terminal Region) followed by a WH (WH’) domain and ends with a hemi-WH domain (pWH’) (**Fig 4B, teal**). Interestingly, the hemi-WH (pWH) from UFL1 complements the hemi WH (pWH’) from the C-terminus of UFBP1 to form a complete WH domain (**Fig 4B**), thus providing an explanation to why UFBP1 is required to stabilize UFL1. Further support to this model is provided by our Y2H screen which identifies the region spanning 268-298, i.e. the C-terminal portion of UFBP1 to interact with UFL1. Superposition of the WH domains of UFL1 and UFBP1 reveal them to have identical folds except for a β-strand which is lacking in the composite WH domain (**Fig S4D-E**). The coiled-coil nature of UFBP1 and the helical backbone of UFL1 may explain the unusual migration of UFL1/UFBP1 by SEC. To validate the predicted structure of the UFL1/UFBP1 complex, we deleted the N-terminal helix along with pWH of UFL1 and checked for complex formation with UFBP1. Indeed we failed to obtain any protein upon deletion of this region of UFL1. Likewise, removal of C-terminal pWH domain of UFBP1 also resulted in poor expression, highlighting the importance of the formation of the composite WH (pWH-pWH’) domain for complex formation and protein stability.

**Figure 4:**
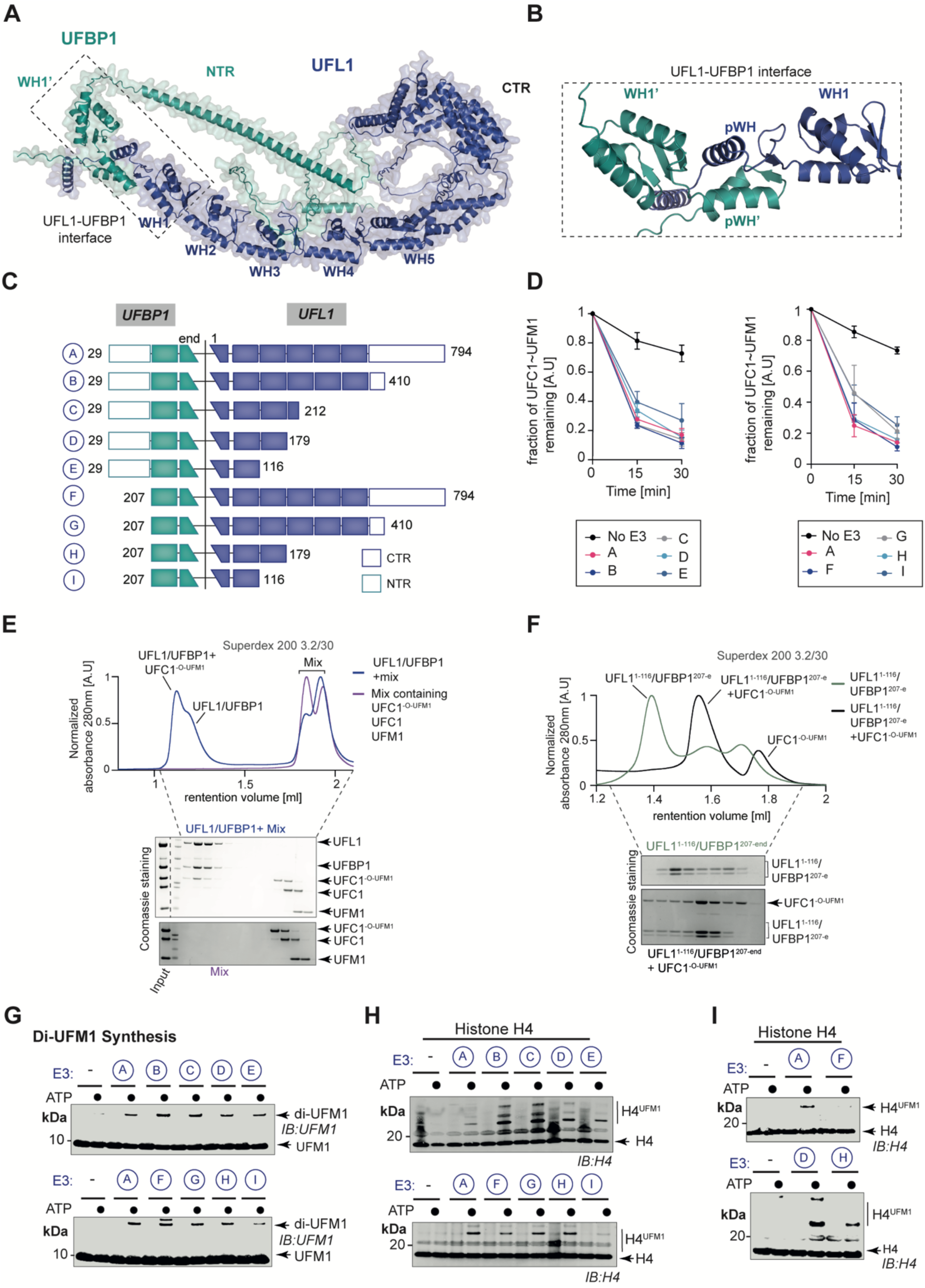
UFL1/UFBP1 complex functions as a scaffold type E3 ligase. **A.** Cartoon representation of predicted full length human UFL1/UFBP1 complex using Alphafold. UFL1 and UFBP1 are shown in blue and teal colors respectively. The dimeric interface of UFL1 and UFBP1 is highlighted in the dotted box. WH: Winged helix; NTR: N-terminal region; CTR: C-terminal region **B.** Enlarged view of the dimeric interface of UFL1/UFBP1 complex shown to highlight the formation of full WH domain bridged by two partial winged helix domains of UFL1 (blue) and UFBP1 (teal). **C.** Schematic representation of UFL1/UFBP1 constructs with different domain boundaries to identify the minimal catalytic region of UFL1/UFBP1 complex required for aminolysis. **D.** Quantitative representation of Lysine discharge assays to identify the minimal boundaries of UFL1 and UFBP1 required for activation of UFC1 (n=3, mean±SD). **E.** SEC elution profiles showing that full length UFL1/UFBP1 complex interacts specifically with charged E2. UFL1/UFBP1 complex was incubated with UFC1^-O-UFM1^ at a 1:1 molar ratio for 20 min at 4°C and loaded on a Superdex 200 Increase 3.2/300 column. The fractions corresponding to each peak were collected and separated on a 4-12% SDS PAGE gel followed by Coomassie staining. **F.** Minimal catalytic region is sufficient for interaction with charged UFC1. UFL1^(1–, 179)^/UFBP1^(1-116)^ complex was incubated with UFC1^-O-UFM1^ at the molar ratio of 1:1 for 20 min at 4°C and analysed by analytical size exclusion chromatography as described in E). **G.** *In vitro* UFMylation assay to monitor formation of free di-UFM1 chains in the presence of UFL1/UFBP1 complexes bearing different domain boundaries. **H.** Substrate UFMylations assays to check for UFMylation of purified Histone H4 in the presence of different UFL1/UFBP1 truncations. **I.** Role of UFBP1 in substrate UFMylation. *(Top)* Comparison of UFMylation activities of UFL1/UFBP1^29-end^ and UFL1/UFBP1^207-end^ using Histone H4. *(Bottom)* Comparison of UFMylation activities of UFL1^1-179^/UFBP1^29-end^ and UFL1^1-179^/UFBP1^207-end^ using Histone H4.

Guided by the insights from the Alphafold predictions, we made additional truncations to map the minimal catalytic domain of the ligase complex (**Fig 4C**). The truncated complexes expressed well and were purified to homogeneity (**Fig S3F**). We first analysed the ability of the different truncated complexes to promote discharge of UFM1 from UFC1∼UFM1 onto Lys. This revealed that all the truncations tested could discharge UFM1 and the smallest region able to promote discharge contains one WH domain each from UFL1 and UFBP1 and the composite WH domain (**Fig 4D, S5A-B**).

We then monitored the ability of the different UFL1/UFBP1 complexes to bind to UFC1 for which we performed analytical SEC of the different UFL1/UFBP1 constructs with UFC1 on its own or a stable non-reactive UFC1^-O-UFM1^ conjugate where UFM1 is linked to C116S via an oxyester bond (**Fig S5 C-D**). UFC1^-O-UFM1^ is more stable compared to thioster-linked UFC1∼UFM1 and therefore used for these analyses. Post preparation, we incubated UFC1^-O-UFM1^ with UFSP2 or 0.2N NaOH and observed complete collapse only in alkaline conditions confirming that the purified UFC1^-O-UFM1^ is indeed linked via an oxyester linkage **(Fig S5E)**. Analysis of complex formation revealed that UFL1/UFBP1 does not bind to UFC1 or UFM1 on its own but forms a stable complex with UFC1^-O-UFM1^ that is of high enough affinity to elute as a complex on SEC (**Fig 4E**). Interestingly, even the smallest of the UFL1/UFBP1 variants was able to bind to UFC1^-O-UFM1^ (**Fig 4F**). Taken together, the ligase complex containing the N-terminal segment of UFL1 (pWH-WH1) and the C-terminal region of UFBP1(WH1’-pWH’), which we refer to as UFL1/UFBP1^min^ (denoted as complex I in **Fig 4C**) is sufficient for binding to charged UFC1 and may represent the minimal ligase domain sufficient to activate UFC1∼UFM1 for aminolysis.

Next, we assayed ligase activity using UFMylation assays monitoring diUFM1 formation. Like full length UFL1, the different C-terminal truncations co-expressed with UFBP1 did not exhibit any loss of activity (**Fig 4G**). Thus, the N-terminal region of UFL1 containing (pWH-WH1) is sufficient for its ligase activity. Similarly, truncation of the N-terminus region (NTR) of UFBP1 did not affect diUFM1 formation. This suggests that the minimal ligase domain is sufficient to form diUFM1 chains. Lastly, we checked the impact of truncations of different regions of UFL1/UFBP1 complex on their ability to modify the substrates such as Histone H4 and MRE11 (**Fig 4G, S5F**). Intriguingly, while deletion of regions outside of WH1, WH2 did not alter diUFM1 formation, these deletions affected substrate modification. Further, deletion of the N-terminal region (NTR) of UFBP1 significantly impacted substrate UFMylation suggesting roles for these regions of the ligase complex in substrate recognition (**Fig 4H-I**). Hence, using systematic truncations based on AlphaFold predictions, we here make the surprising discovery that three tandem WH domains **(Fig S4G)** of UFL1/UFBP1 mediate E3 ligase activity.

### CDK5RAP3 restricts the E3 ligase activity of UFL1/UFBP1

Our Y2H screen identified a second hit with low significance, CDK5 regulatory subunit-associated protein 3 (CDK5RAP3, also known as LZAP), an evolutionarily conserved 53 kDa protein that lacks any known functional domains or motifs. Secondary structure analysis predicts CDK5RAP3 to contain a central unstructured region flanked by *α*-helical stretches at the N and C terminus (**Fig S6**). To determine if CDK5RAP3 can interact with UFL1 in the context of the UFL1/UFBP1 complex, we incubated recombinant full length CDK5RAP3 with UFL1/UFBP1 and analysed complex formation by SEC. Analysis of UV chromatograms and the fractions obtained, confirmed that CDK5RAP3 interacts with UFL1/UFBP1 *in vitro* and forms a stable complex (**Fig 5A**). We further analysed the complex by mass photometry which confirmed the presence of a stable ternary complex with an experimental molecular mass of 192 kDa (**Fig 5B**).

**Figure 5:**
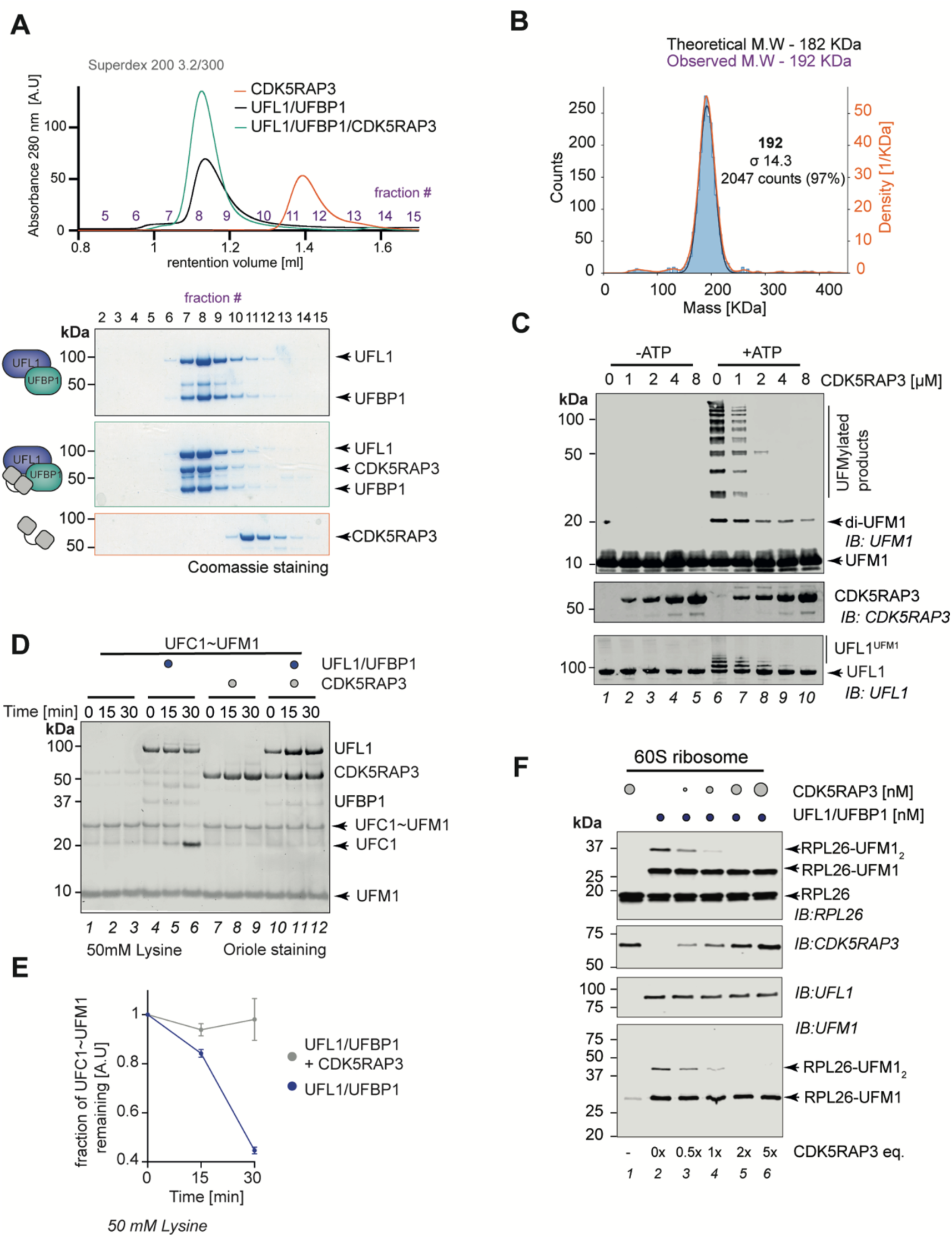
CDK5RAP3 inhibits E3 ligase activity of UFL1/UFBP1 *in vitro*. **A.** CDK5RAP3 forms a complex with UFL1/UFBP1 *in vitro*. 30 *µ*g of UFL1/UFBP1 was mixed with 15 *µ*g of CDK5RAP3 and loaded on an analytical gel filtration column. (Bottom) Fractions were collected and analysed on 4-12% SDS PAGE and visualized by Coomassie staining. **B.** Mass photometry analysis showing the experimental molecular weight of UFL1/UFBP1/CDK5RAP3 complex. **C.** *In vitro* UFMylation assay in the presence of increasing concentrations of CDK5RAP3 to monitor the E3 ligase activity of UFL1/UFBP1 complex. The reaction products were run on a 4-12% SDS-PAGE gel and immunoblotting was performed using indicated antibodies. **D.** Lysine discharge assays to check for activation of UFC1 by UFL1/UFBP1 in the presence and absence of CDK5RAP3. The reaction products were run on a 4-12% SDS PAGE and visualized by Oriole staining. **E.** Quantification of discharge of UFM1 from UFC1 in the presence and absence of CDK5RAP3 as seen in E). n=3, mean ±SD. **F.** Substrate UFMylation assays using purified 60S Ribosomes in the presence of increasing concentration of CDK5RAP3. Reaction was performed for 10 min, stopped by addition of SDS Loading dye and analysed on an SDS-PAGE gel under reducing conditions followed by Immunobloting using indicated antibodies.

Since CDK5RAP3 forms an integral complex with UFL1/UFBP1 and CDK5RAP3 has recently been suggested to function as a substrate adaptor ^53^, we wondered if it could influence E3 ligase activity or substrate UFMylation . We therefore performed *in vitro* UFMylation assays and monitored the E3 ligase activity of UFL1/UFBP1 in the presence of increasing concentrations of CDK5RAP3. Surprisingly, incubation of CDK5RAP3 with UFL1/UFBP1 impaired E3 ligase activity and UFMylation in a concentration dependent manner (**Fig 5C**). Moreover, preassembled UFL1/UFBP1/CDK5RAP3 complexes purified by gel filtration also showed no ligase activity (**Fig S7A**). These observations imply that CDK5RAP3 binds to inhibit the ligase complex.

To gain insights into how CDK5RAP3 inhibits UFMylation, we tested if CDK5RAP3 inhibits complex formation between UFL1/UFBP1 and UFC1^-O-UFM1^. When analysed by pull down and analytical SEC, we find that CDK5RAP3 forms a complex together with UFL1/UFBP1 and UFC1^-O-UFM1^ (**Fig S7B-C**). Since CDK5RAP3 does not affect UFC1∼UFM1 binding, we next tested if CDK5RAP3 binding influences discharge of UFM1 from UFC1∼UFM1 by monitoring transfer of UFM1 onto free Lys. While UFM1 is readily discharged onto Lys in the presence of UFL1/UFBP1, this is completely blocked in the presence of CDK5RAP3 (**Fig 5D-E, S7D**). These observations suggest that binding of CDK5RAP3 to UFL1/UFBP1/UFC1∼UFM1 complex may prevent activation of UFC1∼UFM1 resulting in inhibition of UFMylation.

While these *in vitro* experiments clearly demonstrate that CDK5RAP3 blocks UFC1 activation and UFMylation, it raises the question of the role of CDK5RAP3 in substrate UFMylation. Therefore, we performed UFMylation assays to check for modification of H4, MRE11 and TRIP4 and observed that addition of increasing concentrations of CDK5RAP3 decreased UFMylation of these substrates **(Fig S7E)**. Next, we decided to check the role of CDK5RAP3 in UFMylation of ribosomes, the best described UFMylation substrate to date ^5,6^. Hence, we set up a cell-free reconstitution of ribosome UFMylation. Strikingly, addition of UFL1/UFBP1 to purified 60S ribosomes leads to robust modification of RPL26 (**Fig 5F**), and in line with previous observations, both mono- and di-UFMylated RPL26 are observed. When increasing concentrations of CDK5RAP3 is titrated into this UFMylation reaction, the di-UFMylation of RPL26 is completely abolished (**Fig 5F**). Importantly, even at the highest concentration of CDK5RAP3, monoUFMylation of RPL26 is not affected. Hence, we propose CDK5RAP3 to be a specificity determinant, directing activity of the ligase complex towards substrate Lys by limiting spurious and off-target UFMylation.

### Regulation of UFC1 activity by its N-terminal helix extension

In addition to its core UBC fold, UFC1 has at its N-terminus a highly conserved *α*-helix (*α*0) whose function is unknown (**Fig 6A**). Moreover, UFC1 lacks the oxyanion hole stabilizing Asn as part of the highly conserved HPN motif found in E2 enzymes, which is instead replaced by a TAK motif at this position (**Fig 6B**). Further motivated by the fact that previous studies have shown that the N- and C-terminal extensions on E2s regulate E2 activity ^27^, we sought to determine if *α*0 has a role in regulating UFMylation. First, we first compared the activity of UFC1^WT^ and UFC1 lacking *α*0 (UFC1^ΔN^) in UFMylation assays containing UFC1 with or without UFL1/UFBP1. Surprisingly, UFC1^ΔN^ showed stronger overall UFMylation compared to UFC1^WT^ in the presence of UFL1/UFBP1 (**Fig 6C**). This suggests an inhibitory role for the N-terminal helical extension of UFC1. Next, we compared discharge of UFM1 from UFC1∼UFM1 and UFC1^ΔN^∼UFM1 onto free Lys. In the absence of E3 ligase, UFC1^ΔN^ rapidly discharges UFM1 onto Lys (**Fig 6D & E**) suggesting an increase in its intrinsic lysine reactivity. In the presence of UFL1/UFBP1, discharge of UFM1 onto Lys is enhanced when *α*0 of UFC1 is deleted (**Fig 6D & F**). Taken together with the increase in UFMylation in the presence of UFL1/UFBP1, these results further cement an inhibitory role for the N-terminal helix of UFC1.

**Figure 6:**
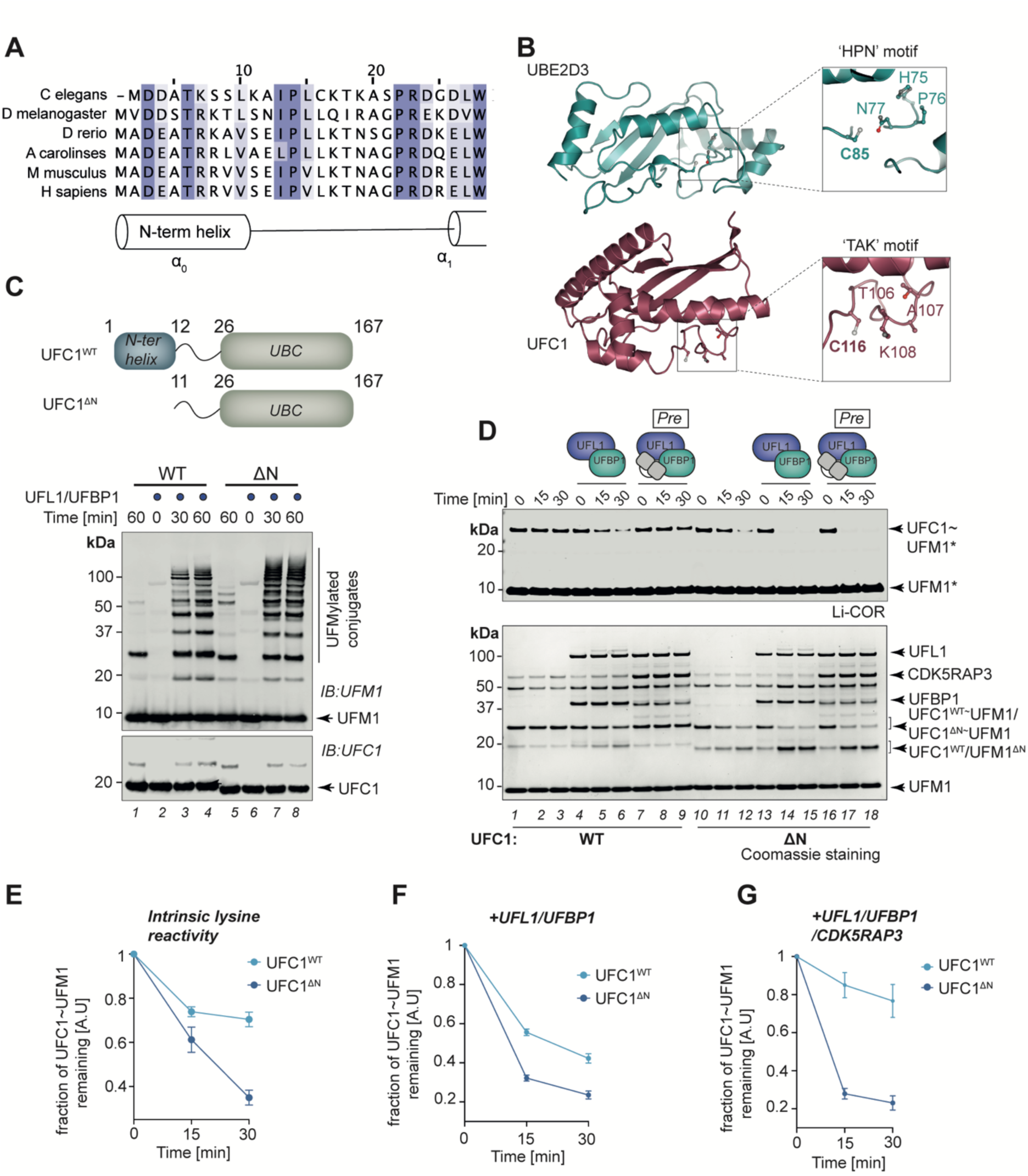
Role of the N-terminal helix of UFC1 in UFMylation. **A.** Multiple sequence alignment of UFC1 homologs from various organisms. A graphical representation of the secondary structure is extracted from the crystal structure of UFC1 (PDB ID:2Z6O) **B.** Crystal structures of UBE2D3 (PDB ID:5EGG) and UFC1 (PDB ID:2Z6O) are shown in cartoon representation. *(Inset)* Catalytic cysteines and critical residues required for catalytic activity of UBE2D3 (green) and UFC1(dark red) are labelled and shown in ball and stick representation. **C.** *In vitro* UFMylation assays in the presence of Wild type UFC1 (UFC1^WT^) and UFC1 lacking the N-terminal helix (UFC1^ΔN^). Schematic representation of domain features of UFC1^WT^ and UFC1^ΔN^ is shown above. **D.** Assay to compare Lys discharge activities of UFC1^WT^ and UFC1^ΔN^ on its own, in the presence of UFL1/UFBP1 or in the presence of preassembled UFL1/UFBP1/CDK5RAP3 complex. Top gel: LiCOR scan of fluorescently labelled UFM1(UFM1*); bottom gel – Coomassie stained (Representative of three independent experiments). **E.** Quantitative analysis of the intrinsic Lys reactivity (Lysine 25 mM) of UFC1^WT^ and in UFC1^ΔN^ in the absence of E3 ligase. **F.** Quantitative analysis of Lys reactivity of UFC1^WT^and in UFC1^ΔN^ in the presence of UFL1/UFBP1. **G.** Quantitative analysis of Lys reactivity of UFC1^WT^and in UFC1^ΔN^ in the the presence of preassembled UFL1/UFBP1/CDK5RAP3. E – G: n=3, mean ±SD.

Since we identified the N-terminal *α*0 of UFC1 to restrain UFMylation, we analysed if inhibition by CDK5RAP3 was mediated via this helix. Indeed, UFMylation assays together with discharge assays comparing activity of UFC1^WT^ and UFC1^ΔN^ reveal that the inhibition of E3 ligase activity by CDK5RAP3 requires the N-terminal helix of UFC1 (**Fig 6D & G, S8**). Based on these results we propose that CDK5RAP3 may interact with both UFL1/UFBP1 and *α*0 of UFC1 to clamp the complex in an inhibited state. Together these analyses reveal a previously unappreciated regulatory role for the N-terminal helix of UFC1 in modulating UFMylation.

## Discussion

In this study, we establish a robust *in vitro* reconstitution system using purified active components of the UFM1 enzymatic pathway to reveal the minimal requirements for UFMylation and mechanistic insights into the ligase machinery. Previous reports have provided conflicting views on the roles of UFBP1 and CDK5RAP3, with some suggesting that they are mainly substrates of UFMylation ^13,29^. Further, it has been suggested that UFBP1 must be UFMylated first at K267 before it can associate with UFL1 to subsequently support UFL1 ligase activity ^12^. Since both UFL1 and UFBP1 complexes are purified from bacteria, our work unequivocally demonstrates that UFL1 and UFBP1 can associate in the absence of any PTMs and the formation of this complex is essential for it to function as an E3 ligase.

It has long been debated as to whether UFL1 is a scaffold/adaptor-type E3 ligase or a Cys-based HECT-like enzyme. Scaffolding type E3 ligases such as RING E3 ligases bind to both E2 and substrate to bring them together for substrate transfer. Since UFL1 can bind to both UFC1 and its supposed substrate UFBP1 at the same time, it was suggested that UFL1 maybe a scaffold-type E3 ligase ^3^. More recently, a UFMylation assay that relied on UFL1 present in mammalian cell extracts to which *in vitro* generated biotinylated E2∼UFM1 thioesters was added, found that UFMylation occurred even when cell extracts were treated with cysteine-alkylating reagents, further suggesting that UFL1 could be a scaffold-type E3 ligase ^13^. Our mutational analysis showing that the UFL1/UFBP1 complex lacks a conserved catalytic Cys combined with the ability of the E3 ligase to promote UFC1∼UFM1 aminolysis establish UFL1/UFBP1 as a scaffold-type E3 ligase. Further reinforcing this conclusion is the observation that K69-linkage specificity is imparted by the E2 in ligase free di-UFM1 formation assays and this K69-linkage specificity is maintained by the E3 ligase. Indeed, this is a feature observed in RING E3 ligases where linkage-specificity is determined by the E2 enzyme ^31,34^. Since UFL1/UFBP1 promotes aminolysis, we speculate that UFL1/UFBP1 binding induces a closed UFC1∼UFM1 conformation akin to RING and atypical SUMO E3 ligases ^54–58^.

UFC1 differs from prototypical E2s since it lacks the catalytic HPN motif, has an exposed catalytic Cys with a lower pKa, lacks the C-terminal *α*-helix observed in canonical E2s and has an additional *α*-helix at its N-terminus (*α*0) ^25,40,59^. Further, it is unclear if the canonical “backside” interaction can occur in UFC1 ^28,60^. These unique features of UFC1 and the unconventional features of the ligase complex makes it likely that the UFM1 machinery employs a unique mechanism to transfer UFM1 from UFC1 onto substrate. Interestingly, the UBE2E enzymes also have an intrinsically disordered N-terminal extension that have an inhibitory role restricting Ub transfer thereby limiting polyUb chain formation ^61^. Future work will reveal whether E3 binding induces a closed UFM1∼UFM1 conformation and whether *α*0 impedes this process. Nevertheless, our data seem to suggest that *α*0 has a regulatory role potentially preventing UFM1 discharge in the absence of the E3 ligase.

The recent advances in structure prediction ^50^ enabled us to narrow the regions of UFL1/UFBP1 essential for ligase activity to a tandem repeat of three WH domains - a WH domain each from UFL1 and UFBP1 and the composite WH domain formed by the two proteins. As this region can bind to UFC1∼UFM1 and activate the E2 for UFM1 discharge, we define this to be the minimal catalytic domain. Further structural studies will be required to reveal why three WH domains are required to form a functional ligase complex and the identification of linchpin residues in UFL1/UFBP1 that play similar roles to RING E3 ligases to activate the E2 for aminolysis ^55,56^. Intriguingly, the anaphase promoting complex (APC) subunit APC2 contains a WHB domain that binds the “backside” of UCBH10/UBE2C, located at a face opposite from the catalytic site and is an allosteric site in several E2s ^62,63^. This backside binding of the WHB domain is specific towards UBE2C and is important for transfer of Ub onto specific substrates ^63^. Our observations also raise the question of the function of the other WH/PCI domains in UFL1/UFBP1 to ligase function. One possibility is that the WH domains may mediate substrate recognition, binding to UFM1 or serve to either regulate recruitment or processivity of chain formation. Given that the WH domain of Cockayne syndrome group B (CSB) can bind to ubiquitin, it is tempting to speculate that one of the WH domains of UFL1/UFBP1 can bind to UFM1 ^64^. WH domains commonly recognize nucleic acids and are typically found in transcription factors or as protein interaction domains ^65^. To our knowledge, this is the first demonstration of catalytic activity mediated by WH domains ^66^, leading us to propose moonlighting functions for other WH domain containing proteins. It also raises the possibility that other WH domains could have E3 ligase activity for UFMylation or other UBLs. Since UFL1 has been proposed to have nuclear functions in telomere maintenance and DNA damage response ^10,11,67^, one possibility is that the WH domains may mediate DNA binding. A further possibility is the use of the WH domains of UFL1/UFBP1 to recognize rRNA or translating mRNA during ribosome UFMylation.

Our *in vitro* studies show that formation of UFM1 chains and autoUFMylation is inhibited by CDK5RAP3 (**Fig 2C**). Indeed, previous studies have observed an altered pattern of UFMylation in the absence of CDK5RAP3 leading to the suggestion that it may be a substrate adaptor ^53^. Interestingly, CDK5RAP3 was suggested to function as a sensor for ER stress to induce autophagic degradation of aberrant proteins formed as a result of ribosomal stalling ^68^. Several multidomain and multiprotein E3 ligases such as PARKIN and Cullin Ring Ligases (CRLs) are inhibited and their activation is a carefully orchestrated multistep process ^35,69^. Based on our observations, we propose that ligase complexes containing UFL1, UFBP1 and CDK5RAP3 represent an autoinhibited state. The surprising finding that ribosome UFMylation is not abolished in the presence of CDK5RAP3 but rather restricted to monoUFMylation leads us to suggest that CDK5RAP3 regulates ligase activity in the following manner: (i) in the absence of substrate, binding of CDK5RAP3 inhibits E3 ligase activity; (ii) this autoinhibition is relieved when substrates such as the 60S ribosome are encountered. This release from inhibition may involve conformational changes induced upon recognition of structural features on the substrate by UFL1, UFBP1 or CDK5RAP3 to mediate substrate UFMylation. Such multi-layered regulation possibly prevents spurious UFMylation and ensures ribosome UFMylation only in the right context. While previous studies have suggested roles for UFMylation for UFL1-UFBP1 interaction, and UFMylation and phosphorylation to enhance UFMylation ^10,12,19,29^, our minimal reconstitution clearly demonstrates that the ligase complex assembles in the absence of any PTM. Although we cannot rule out a role for phosphorylation in enhancing ligase activity, the rapid UFMylation of ribosomes by the reconstituted ligase suggests that it may be unlikely.

Our rebuilding approach provides insights into the assembly, minimal requirements, and mechanism of the UFL1/UFBP1/CDK5RAP3 ligase and reveals principles of protein UFMylation. The *in vitro* reconstitution system we describe here using purified components lays the foundation for future biochemical and structural studies to understand the molecular mechanism of this unusual E3 ligase complex and UFMylation. Further, the cell-free UFMylation system can be applied to understand the logic of ribosome UFMylation and its relationship to ribosome quality control pathways.

## Acknowledgements

We thank members of the Kulathu lab for critical reading of the manuscript and helpful comments. We thank the Zeqiraj lab (Univ of Leeds) and Prof Hofmann (Univ of Köln) for helpful discussions and advice. We acknowledge MRC Reagents and Services for providing valuable reagents, especially Drs. Simone Weidlich and Rachel Toth for cloning the constructs used in this study. We thank Mr. Joby Varghese for help with Mass spectrometry data acquisition and preliminary analysis and Dr. Emma Brannigan for help with mass photometry measurements. We thank Christopher Lapointe (Puglisi lab, Stanford) for valuable reagents and discussion. We also thank Dr. Juraj Ahel (IMP Vienna) for his help with generating composite foldindex profile of UFL1. This research was supported by ERC Starting grant (RELYUBL, 677623), the Lister Institute of Preventive Medicine, BBSRC (BB/T008172/1) and MRC grant MC_UU_00018/3 to YK; Grant R01GM074874 from the National Institute of General Medical Sciences and a generous gift from the Becker Family Foundation to R.R.K; NIH Epilepsy Training Program T32 NS007280 to PAD.

## Author contributions

YK and JJP conceived the study. JJP designed, performed and analysed all the experiments. HM designed, expressed and purified UFL1/UFBP1 constructs based on Alphafold. PADR performed *in vitro* ribosome UFMylation assays. DM and SPM contributed reagents and RS performed SEC MALS analysis. YK analysed data, supervised the study and secured funding. JJP and YK wrote the manuscript with input from all the authors.

## Declaration of interests

The authors declare no competing interests.

## Supplemental information

Supplemental information includes Figures S1-S8 and Table S1.

## Materials and Methods

### Plasmids, Cloning and mutagenesis

The details of cDNA constructs used in this study is given in Table S1. Cloning of most of the constructs was performed by MRC PPU Reagents and Services (University of Dundee). Briefly, mutagenesis was carried out using Q5 site-directed mutagenesis kit (NEB) with KOD polymerase (Novagen) according to manufacturer’s protocol. Following mutagenesis, cDNA constructs were amplified using *E. coli* DH5α and purified using QIAprep spin mini-prep kit (Qiagen). All cDNA constructs were verified by DNA sequencing and services, University of Dundee using DYEnamic ET terminator chemistry (Amersham Biosciences) on Applied Biosystems automated DNA sequencers.

### Recombinant protein expression and purification

Recombinant His_6_-3C-UBA5, His_6_-3C-UFM1^WT^, His_6_-3C-UFM1^Konly^, His_6_-3C-UFM1^KtoR^, His_6_-3C-Cys-UFM1, His_6_-3C-UFC1^WT^, His6-3C-UFC1^ΔN^ and His_6_-3C-MRE11 were expressed in *E. coli* BL21 and purified using Ni^2+^-NTA affinity chromatography as the first step. Briefly, *E. coli* BL21 cultures expressing His_6_-tagged proteins were grown in 2xTY medium at 37°C until OD_600_ reached 0.6-0.8. Final concentration of 0.3 mM IPTG was added and the cultures were incubated at 18 °C for 16 h. Cells were harvested, resuspended, and lysed in appropriate lysis buffer containing 25 mM Tris pH 8, 300 mM NaCl, 10% Glycerol, 2 mM DTT, 1 mM Benzamidine, 1 mM AEBSF, 1x protease inhibitor cocktail (Roche) by ultrasonication. Lysed cells were then clarified by centrifugation at 30000g for 30 mins at 4°C. The clarified lysate was then incubated with pre-equilibrated Ni^2+^-NTA Agarose beads (Amintra, Abcam) for 2 h in appropriate binding buffer containing 25 mM Tris pH 8, 300 mM NaCl, 10% Glycerol,10 mM Imidazole. The beads were then washed extensively using wash buffer containing 25 mM Tris pH 8, 300 mM NaCl, 10% Glycerol, 2 mM DTT and 20 mM Imidazole. Finally, the bound protein was eluted using elution buffer containing 25 mM Tris pH 8, 300 mM NaCl, 10% Glycerol, 2 mM DTT and 300 mM Imidazole. Wherever necessary, His_6_-tags were cleaved off by incubating tagged proteins with PreScission protease at 4°C overnight. A final size exclusion chromatography step was performed using Superdex 75 HiLoad^TM^ 16/60 Superdex^TM^ 75pg and HiLoad^TM^ 16/60 Superdex^TM^ 200 pg columns (GE Healthcare Life Sciences) with buffer containing 25 mM Tris pH 8.0, 150 mM NaCl, 10% Glycerol and 2 mM DTT. The purified proteins were then concentrated using Amicon^TM^ Ultra 15 concentrators (MERCK Millipore) and stored in -80°C.

### Purification of GST-tagged proteins

GST-TEV-TRIP4 was expressed in *E. coli* BL21(DE3) strain as described above. Cells were harvested and lysed in lysis buffer containing 25 mM Tris pH 7.5, 300 mM NaCl, 10% Glycerol and 2 mM DTT using ultrasonication. Pre-equilibrated Glutathione 4B-sepharose beads (Amintra, abcam) were incubated with clarified lysate for 2 h. The beads were then washed with high salt buffer containing 25 mM Tris pH 7.5, 500 mM NaCl, 10% glycerol and 2 mM DTT. Further the beads were washed with low salt buffer containing 25 mM Tris pH 8, 150 mM NaCl, 10% glycerol and 2 mM DTT. The protein was then cleaved off the tag by incubation with Precision protease at 4°C overnight. A second ion exchange chromatography step was carried out using Resource Q (6ml) (GE Healthcare Life Sciences) column with low salt buffer as 25mM Tris pH 7.5, 150 mM NaCl, 10% glycerol, 2 mM DTT and high salt buffer as 25mM Tris pH 7.5, 500mM NaCl, 10% glycerol and 2 mM DTT. A final size exclusion chromatography step was performed using HiLoad^TM^ 16/60 Superdex^TM^ 75pg (GE Healthcare Life Sciences) and the protein was buffer exchanged in buffer 25 mM Tris pH 7.5, 150 mM NaCl, 10% glycerol and 2 mM DTT. The purified proteins were then concentrated and stored in -80°C.

### Co-expression and purification of UFL1-UFBP1 complex

His_6_-TEV-UFL1 and StrepII-C3-UFBP1 were cloned in a pET Duet1 construct and expressed in *E. coli* BL21 codon plus (RIPL) (Agilent) cells. Bacterial cultures were grown in 2xTY medium at 37°C until OD_600_ reached 0.6. Final concentration of 0.3 mM IPTG was added to induce the expression and the cultures were incubated at 18°C for 16 h. Cells were harvested, resuspended in buffer containing 25 mM Tris pH 8.0, 300 mM NaCl, 2 mM DTT, 1mM Benzamidine, 1mM AEBSF, protease inhibitor cocktails (Roche). Cells were lysed by High pressure homogenization using French press (Avestin Emulsiflux C3, Wolflabs). The lysate was then clarified by Ultracentrifugation at 30000g for 30 mins at 4°C. Affinity purification was carried out in two steps. In the first step, the clarified lysate (25 mM Tris pH 8.0, 300 mM NaCl, 20 mM Imidazole, 2 mM DTT) was passed through HisTrap^TM^ FF (GE Healthcare Life Sciences) column. After binding, the column was washed with 10 column volumes (CV) of binding buffer containing 25 mM Tris pH 8.0, 300 mM NaCl, 20 mM Imidazole, 2 mM DTT. After washing, bound proteins were eluted using buffer containing 25 mM Tris pH 8.0, 300 mM NaCl, 300 mM Imidazole, 2 mM DTT by applying a concentration gradient of Imidazole. The eluted proteins were then passed through StrepTrap^TM^ (GE Healthcare Life Sciences) column pre-equilibrated with buffer containing 25 mM Tris pH 8.0, 300 mM NaCl, 2 mM DTT. After 2 CV of washing with binding buffer, the proteins were eluted using 25 mM Tris pH 8.0, 300 mM NaCl, 2 mM DTT and 2.5 mM Desthiobiotin. Finally, purified proteins were passed through HiLoad^TM^ 16/60 Superdex^TM^ 200pg (GE Healthcare Life Sciences). The purified protein was then stored in -80°C in buffer containing 25 mM Tris pH 8.0, 300 mM NaCl, 5% Glycerol and 2 mM DTT until further use.

### Preparation and labelling of IRDye800CW-UFM1

Cys-UFM1^1–83^ which contains a Cys residue upstream of M1 was purified and exchanged into a fresh buffer containing 50 mM HEPES pH 7.5, 150 mM NaCl and 0.5 mM TCEP using CentriPure P10 columns (EMP Biotech). Protein was then mixed with IRDye^®^ 800CW Maleimide (LI-COR^®^) at 5:1 molar ratio and incubated for 2 h at RT. Unreacted dye was removed by three-step buffer exchange process using CentriPure P10 columns (EMP Biotech) (once) and PD-10 Sephadex G-25 column (twice) in a sequential manner according to manufacturer’s instruction. Finally, to remove residual unreacted dye material, the labelled protein was dialyzed into buffer containing 25 mM Tris pH 8, 150 mM NaCl and 2 mM DTT using dialysis membrane (Thermo Scientific) (MWCO 3000) overnight at 4°C and stored at -80°C until further use.

### Biochemical assays

#### Lysine discharge assays

To analyse the intrinsic reactivity of UFC1, a substrate-independent single turnover discharge assay was employed. Firstly, UFC1 was charged with UFM1^WT^ or labelled UFM1 by incubating 0.5 *µ*M UBA5, 10 *µ*M UFC1 and 20 *µ*M UFM1 in a buffer containing 50 mM HEPES pH 7.5, 50 mM NaCl, 0.5 mM DTT and10 mM MgCl_2_. The charging reaction was initiated by addition of 10 mM ATP and incubation of the reaction mix at 37°C for 20-30 mins. To stop further charging of UFM1 onto UFC1, the reaction was quenched by addition of 50 mM EDTA and subsequent incubation at RT for 10 mins. To check for discharge, the quenched mixture was incubated with 50 mM lysine (pH 8.0) or other amino acids namely Serine, Threonine, Arginine, Cysteine at 37°C. The reaction was stopped at each of the time points by addition of SDS-Loading dye without any reducing agent to the reaction mix and run on a 4-12% SDS-PAGE under non-reducing conditions followed by Coomassie staining or visualization using LI-COR^®^ Odyssey.

In reactions involving the UBE2D3 and UBE2L3, 0.5 *µ*M UBE1 was incubated with 10 *µ*M UBE2D3/UBE2L3 and 20 *µ*M Ub in reaction buffer containing 50 mM HEPES pH 7.5, 50 mM NaCl,10 mM ATP and 10 mM MgCl_2_ at 37°C for 20 mins. The reaction was quenched, and discharge was analysed as described above.

In discharge assays involving UFL1/UFBP1 and CDK5RAP3, the quenched reaction mix was added to a cocktail containing 1 μM UFL1/UFBP1 or 1 *µ*M or 2 *µ*M of CDK5RAP3. Discharge was initiated by addition of 50 mM lysine (pH 8.0) in buffer containing 50 mM HEPES pH 7.5, 50 mM NaCl and 0.5 mM DTT and incubated at 37°C for indicated time duration. The reaction was stopped and analysed as described above.

#### *In vitro* UFMylation assays

To check for UFMylation *in vitro,* 0.25 *µ*M UBA5, 5 μM UFC1, 1 μM UFL1 and 20-30 μM UFM1 were incubated in reaction buffer containing 50 mM HEPES 7.5, 10 mM MgCl_2_ and 5 mM ATP for 1 h at 37°C. The reaction was stopped by the addition of SDS loading buffer containing a reducing agent. The reaction products were then separated on a 4-12% SDS-PAGE gel under reducing conditions and analysed by immunoblotting using indicated antibodies. In UFMylation assays involving substrates, around 1 μM-2 μM of recombinant substrates were incubated with 0.25 μM UBA5, 5 μM UFC1, 1μM UFL1 and 30 μM of UFM1 in buffer containing 50 mM HEPES 7.5,10 mM MgCl_2_ and 5 mM ATP for 1 h at 37°C. Following incubation, the reaction was stopped and analyzed using SDS-PAGE or Immunoblotting using indicated antibodies.

#### Pulldown assays

To analyse the interaction of the E3 ligase with the E2, pulldown assays were performed. Approximately 10 *µ*M of untagged UFC1 or UFC1^-O-UFM1^ was incubated with 5 *µ*M of full length UFL1/UFBP1 in binding buffer containing 25 mM Tris pH 8.0, 300 mM NaCl, 2 mM DTT for 1 h at 4°C. Following incubation, 30 *µ*l of pre-equilibrated Streptavidin Sepharose beads (IBA Life Sciences) (50% slurry) was added and further incubated for about 1h at 4°C. Following binding, centrifuge tubes containing the reaction mix was spun down at 500g for 3 mins and the supernatant was discarded to remove unbound proteins. To further remove weakly bound proteins, the beads were washed thrice with 1 ml of ice-cold binding buffer. Finally, the bound proteins were eluted by addition of binding buffer containing 2.5 mM Desthiobiotin (pH 8.0) and incubation at 4°C for 30 min. The eluates were then analysed on 4-12% SDS PAGE under reducing conditions followed by Coomassie staining.

### Analytical gel filtration chromatography analysis

For analysing the interaction between UFL1/UFBP1 and CDK5RAP3, around 30 *µ*g of UFL1/UFBP1 and 15 *µ*g of CDK5RAP3 were mixed and incubated for 30 min on ice. Then, around 50 *µ*ls of sample was loaded on Superdex^TM^ 200 Increase 3.2/300 column (GE Healthcare Lifesciences) pre-equilibrated with buffer containing 25 mM Tris pH 8.0, 300 mM NaCl and 2 mM DTT. The fractions corresponding to each peak were collected and further analysed on a 4-12% SDS PAGE under reducing conditions followed by Coomassie staining.

To check if UFL1/ UFBP1 could be reconstituted *in trans*, around 15 *µ*g of purified His_6_-UFL1 and StrepII-UFBP1 were mixed and incubated on ice for 2h. After incubation, sample was loaded on Superdex^TM^ 200 Increase 3.2/300 column (GE Healthcare Lifesciences) column pre-equilibrated with buffer containing 25 mM Tris pH 8.0, 300 mM NaCl, 2 mM DTT and the chromatograms were analysed.

### Preparation of UFC1^-O-UFM1^

UFC1^-O-UFM1^ was prepared by incubating 40 *µ*M UFC1^C116S^ with 40 *µ*M UFM1, 1 *µ*M UBA5 in buffer containing 50 mM Tris pH 8.8, 10 mM MgCl_2_, 5 mM ATP overnight at 25°C. The reaction was incubated briefly with 20 mM DTT to remove any non-specific di-sulfide linked adducts formed and passed through HiLoad^TM^ 16/60 Superdex^TM^ 75 pg with buffer containing 25 mM Tris pH 8.0, 150 mM NaCl, 5% Glycerol and 2 mM DTT. The fractions corresponding to UFC1^-O-UFM1^ were collected and analysed on 4-12% SDS PAGE gel. Finally, the fractions containing UFC1^-O-UFM1^ were pooled, concentrated and stored in -80°C until further use.

### Alphafold Predictions

The structure of UFL1/UFBP1 was predicted using the ColabFold Google Colab notebook “AlphaFold_advanced” ^50^ ( https://github.com/sokrypton/ColabFold). The predicted model with the highest IDDT score is shown in the main figure and the PAE scores for different models ranking 1 to 5 are shown in **Fig S4A**.

### DALI Analysis

DALI server was used to perform structural similarity analysis ^70^. WH1 domain of UFL1(52-115) was used as the query model and searched against PDB25 database. The top ten models in the order of the Z-score is shown in **Fig S4B**.

### Bioinformatic analysis

For generation of composite Foldindex profile, previously reported method was adapted ^71,72^. First, individual Foldindex profiles of UFL1 sequence from different organisms were generated locally using a custom foldindex program provided by Juraj Ahel. Then, a composite Foldindex profile was prepared by blending the images in Adobe illustrator with 30% opacity.

For generation of multiple sequence aligments, the protein sequences of target proteins and their homologs were manually downloaded from UNIPROT database in fasta format. Multiple sequence alignment was performed using Jalview version 2.11.1.7 programme (MAFFT algorithm module using L-INS-I) ^73,74^.

### Mass Photometry Data Acquisition and Analysis

Protein samples were prepared in a solution containing 25 mM Tris pH 8.0, 300 mM NaCl, 2 mM DTT and stored on ice before loading onto the Refeyn One^MP^ instrument (Refeyn). Typically, a set of protein standards (NativeMark™ Unstained Protein Standard, Invitrogen) are used for calibration and to generate a standard curve. Following calibration, approximately 10 *µ*l of diluted protein sample in the concentration range of 5–25 nM was introduced into the flow-chamber and movies of 60s duration were recorded. Data acquisition was performed using Acquire MP (Refeyn Ltd, v1.1.3) and analysed using Discover MP software. Data shown here is representation of at least 3 independent acquisitions (*n* ≥ 3).

### SEC MALS data acquisition and analysis

Size exclusion chromatography and multi angle light scattering (SEC–MALS) experiments were performed on a Dionex Ultimate 3000 HPLC system with an inline Wyatt miniDAWN TREOS MALS detector and Optilab T-rEX refractive index detector. In addition, the elution profile of the protein was monitored with UV 280 attached to the Dionex system. For size exclusion chromatography, Superdex 200 10/300 gl column (GE Healthcare LifeSciences) was used. 50 *µ*l of the purified UFL1/UFBP1 stored in buffer containing 25 mM Tris pH 8.0, 300 mM NaCl, 2 mM DTT was loaded into the SEC column with Dionex auto loader at a concentration of 4 mg/ml and a flow rate of 0.3 ml/min was maintained throughout the experiment. Molar masses spanning elution peaks were calculated using ASTRA v6.0.0.108 (Wyatt).

### Ribosome UFMylation assays

Purified 60S ribosomes were a generous gift from the Puglisi lab, and purified as described previously ^75,76^. *In vitro* UFMylation of ribosomes was performed at 30 °C with 1 *µ*M UFC1, 0.5 *µ*M UBA5, 0.5 *µ*M UFM1, 0 nM or 75 nM UFL1/UFBP1, 50 nM purified 60S ribosomes, and increasing concentrations of CDK5RAP3 (0, 38, 75, 150 or 375 nM) in a reaction buffer of 25 mM HEPES pH 7.5, and 100 mM NaCl. The reaction was quenched in 4 x SDS-Load buffer. Western blots show 10 min reaction time; immunoblots for RPL26 (Abcam, 59567) and UFL1 (Bethyl, A303-455M) were run on the same gel, which was cut and probed for these proteins separately. Immunoblots for CDK5RAP3 Bethyl, A300-871A) and UFM1 (Abcam, Ab109307) were run on separate gels.

### LC-MS/MS sample preparation, data acquisition and analysis

First, an *in vitro* UFMylation reaction was performed to generate UFMylated products including free UFM1 chains. Then, the reaction products were run on a 4-12% SDS PAGE gel to separate the products based on electrophoretic mobility. Next, the bands corresponding to di-UFM1 chains were excised and in-gel digestion was performed ^77^. Digested peptides were analysed by LC-MS/MS on an Exploris 480 coupled to an Ultimate 3000 nanoLC system (Thermo Fisher Scientific) for figure 2D and on an Exploris 240 (Thermo Fisher Scientific) coupled to an Evosep One (Evosep) for figure 4D. For the analysis performed on the Ultimate 3000, samples were loaded on a 100 *µ*m x 2 cm trap column (Thermo Fisher Scientific #164564-CMD) and analysed on a 75 *µ*m x 50 cm analytical column (Thermo Fisher Scientific #ES903) using a gradient from 3% to 35% Buffer B (80% LCMS grade acetonitrile, 0.08% formic acid in water) over 53 minutes. The columns were then washed with 95% Buffer B for 2 minutes prior equilibration in 97% Buffer A (0.1% formic acid in LCMS grade water). For the analysis performed on the Evosep One, samples were loaded onto the Evotips as per manufacturer recommendations and analysed using the 30 SPD Method. Peptides were then analysed in on either the Exploris 240 or 480 using data dependant with a MS1 resolution of 60000, AGC target of 300% and maximum injection time of 25 or 28 ms. Peptide were then fragmented using TOP 2 s method, MS2 resolution of 15000, NCE of 30 or 32%, AGC of 100% and maximum injection time of 100 ms. Peptide identification was performed in Mascot using a restricted and frequently updated database containing ∼2000 protein sequences of interest (MRC db). Carbamidomethylation (C) was set at fixed modification and Oxidation (M), Dioxidation (M) and the addition of the dipeptide Glycine-Valine (K) were set as variable modifications. Peptides were searched using an MS1 tolerance of 10 ppm and MS2 tolerance of 0.06 Da and a maximum of 2 missed cleavages were allowed. Only hits identified with a FDR (False Discovery Rate) <1% were selected and further analysed in Scaffold viewer V5. For semi-quantitative analysis of VG modification on individual sites, Total Ion chromatogram (TIC) values obtained from the LC/MS and represented graphically.

---

**Figure S1:**
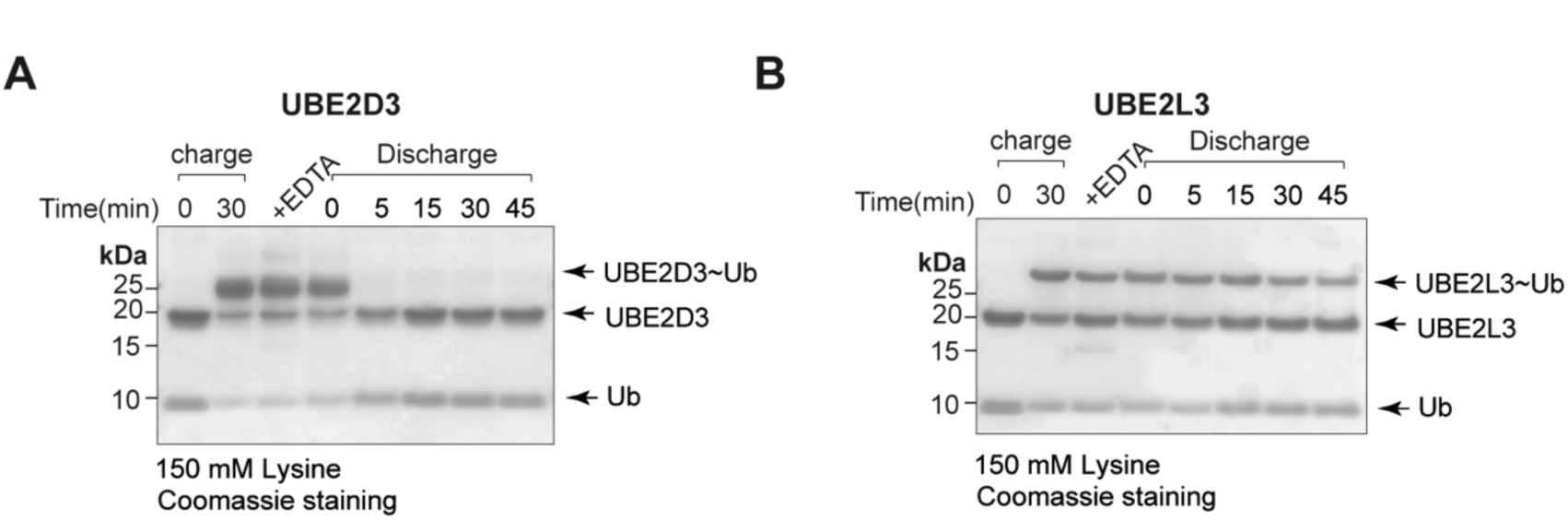
Analysis of E2 discharge. **A.** & **B.** Time-dependent analysis of discharge of Ubiquitin from UBE2D3 and UBE2L3 in the presence of 150 mM.

**Figure S2:**
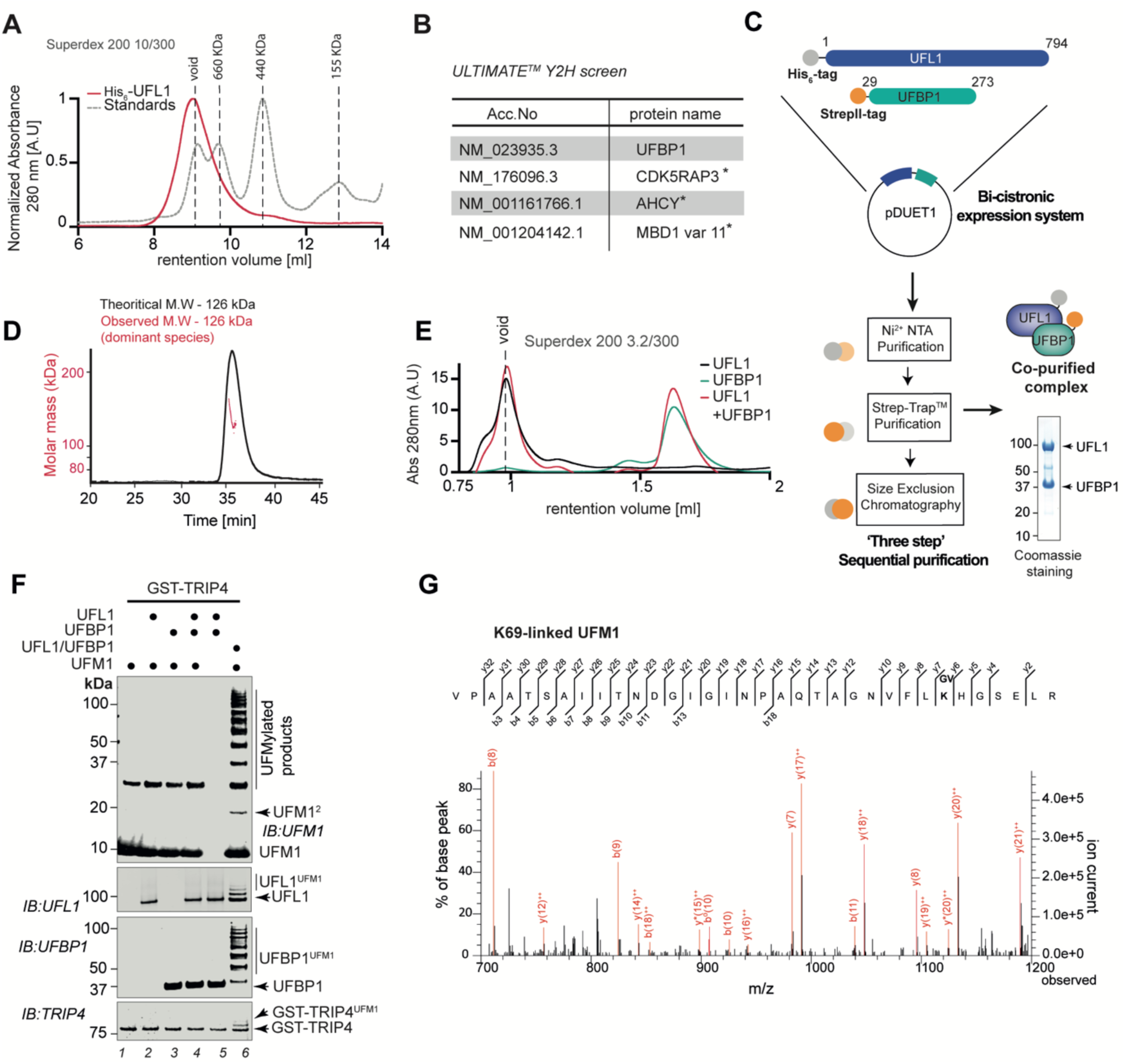
UFL1 is only active in the presence of co-expressed UFBP1. **A.** Size exclusion chromatography profile of UFL1 run on Superdex 200 Increase 10/300gl column (shown in red). Overlay of chromatogram of molecular weight standards of different sizes (shown in grey) run under same buffer conditions. **B.** Results obtained from ULTIMATE Y2H^TM^ screening (Hybrigenics) for binary interactions with UFL1. Asterisk denotes protein hits obtained with low confidence. **C.** Schematic describing the strategy for co-expression and purification of UFL1/UFBP1 complex. **D.** SEC-MALS analysis of UFL1/UFBP1 complex. The theoretical and observed molecular weights are indicated. **E.** Analytical gel filtration chromatography analysis of UFL1/UFBP1 complex and its subunits run on Superdex 200 3.2/300. **F.** UFBP1 does not activate UFL1 when added exogenously. *In vitro* UFMylation assays to compare the E3 ligase activity of UFL1 and UFBP1 expressed alone and together as a complex. 0.25 *µ*M UBA5, 5 *µ*M UFC1 and 10 *µ*M UFM1 was incubated with 1 *µ*M UFL1 or 1 *µ*M UFBP1 or 1 *µ*M UFL1/UFBP1 complex for 1 h at 37°C in the presence of 50 mM HEPES 7.5, 0.5 mM DTT, 10 mM MgCl_2_ and 10 mM ATP. The reaction was stopped by addition of 3x SDS loading dye and run on a 4-12% denaturing SDS PAGE gel under reducing conditions and immunoblotting was performed using indicated antibodies. **G.** MS^2^ spectra showing the peptide derived from *in vitro* UFMylation assay showing the VG-remnant on K69 of UFM1.

**Figure S3:**
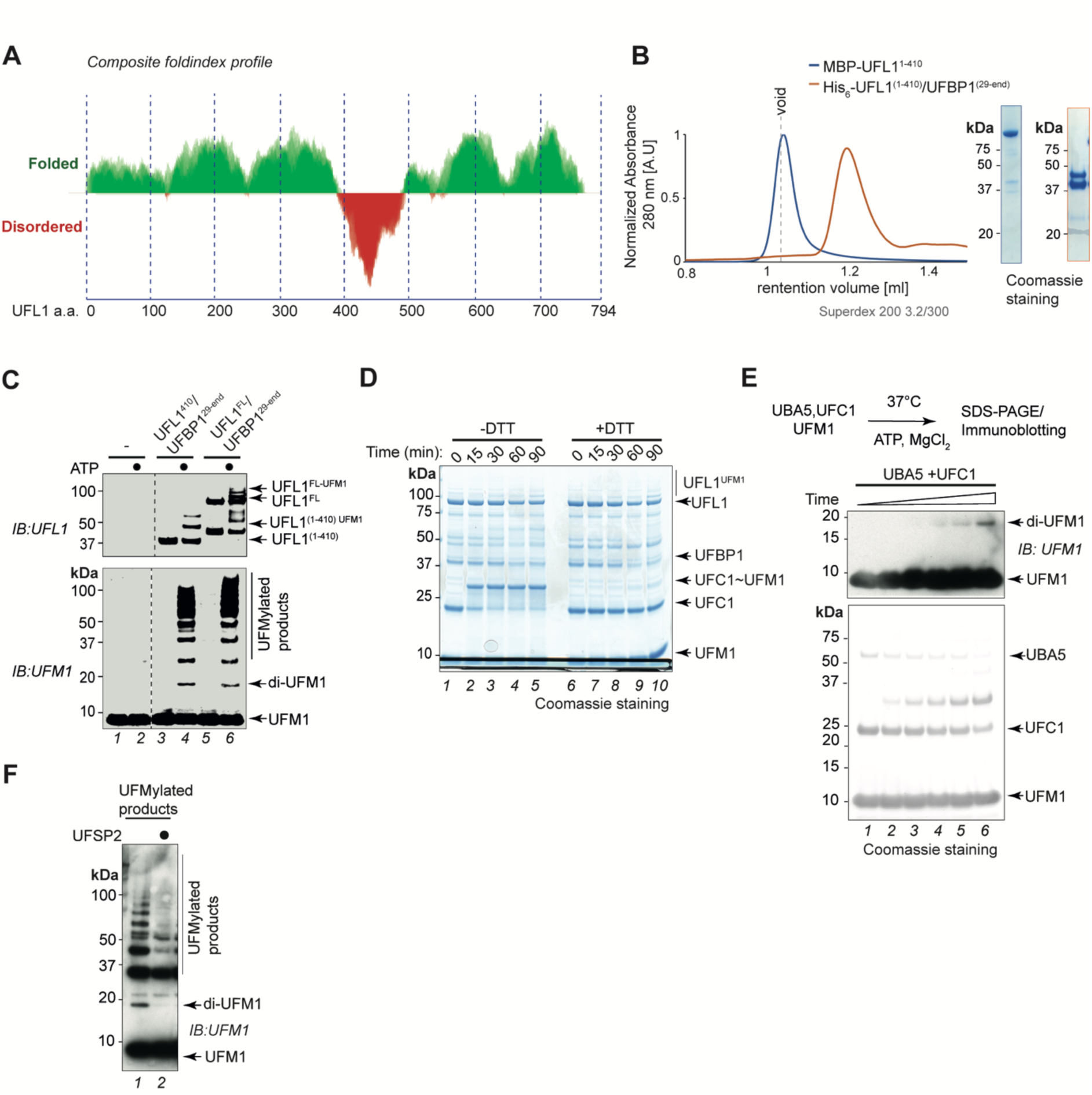
Mode of UFMylation by UFL1/UFBP1 complex. **A.** Composite Foldindex profile of UFL1 showing folding propensity of different regions of UFL1 to aid in construct design for soluble protein expression. **B.** Size exclusion chromatography profile of MBP-UFL1^1-410^ (dark blue) and His_6_-UFL1^1-410^/UFBP1^29-end^(orange) run on Superdex 200 3.2/300 column (shown in red). Approximately 20 *µ*gs of sample was run on a Superdex 200 3.2/300 analytical gel filtration column and UV traces measured at 280 nm were recorded and analysed. (Right) Coomassie stained gel showing the purity of the proteins. **C.** *In vitro* UFMylation assay to check for E3 ligase activity of UFL1^1-410^/UFBP1^29-end^. Full length UFL1/UFBP1 complex is used as a positive control. **D.** Time course assay to check for transthiolation activity of UFL1. Reaction products were analysed on a 4-12% SDS PAGE gel under reducing and non-reducing conditions. **E.** *In vitro* UFMylation assays to check for formation of di-UFM1 chains in the by minimal reconstitution in the presence UBA5 and UFC1 alone. **F.** Immunoblot to check for the presence of UFM1 chains with and without treatment of UFSP2. *In vitro* UFMylation products generated as shown in E was incubated with UFSP2 and analysed by immunoblotting using anti-UFM1 antibody to check for the disspearence of polyUFMylated products especially di-UFM1.

**Figure S4:**
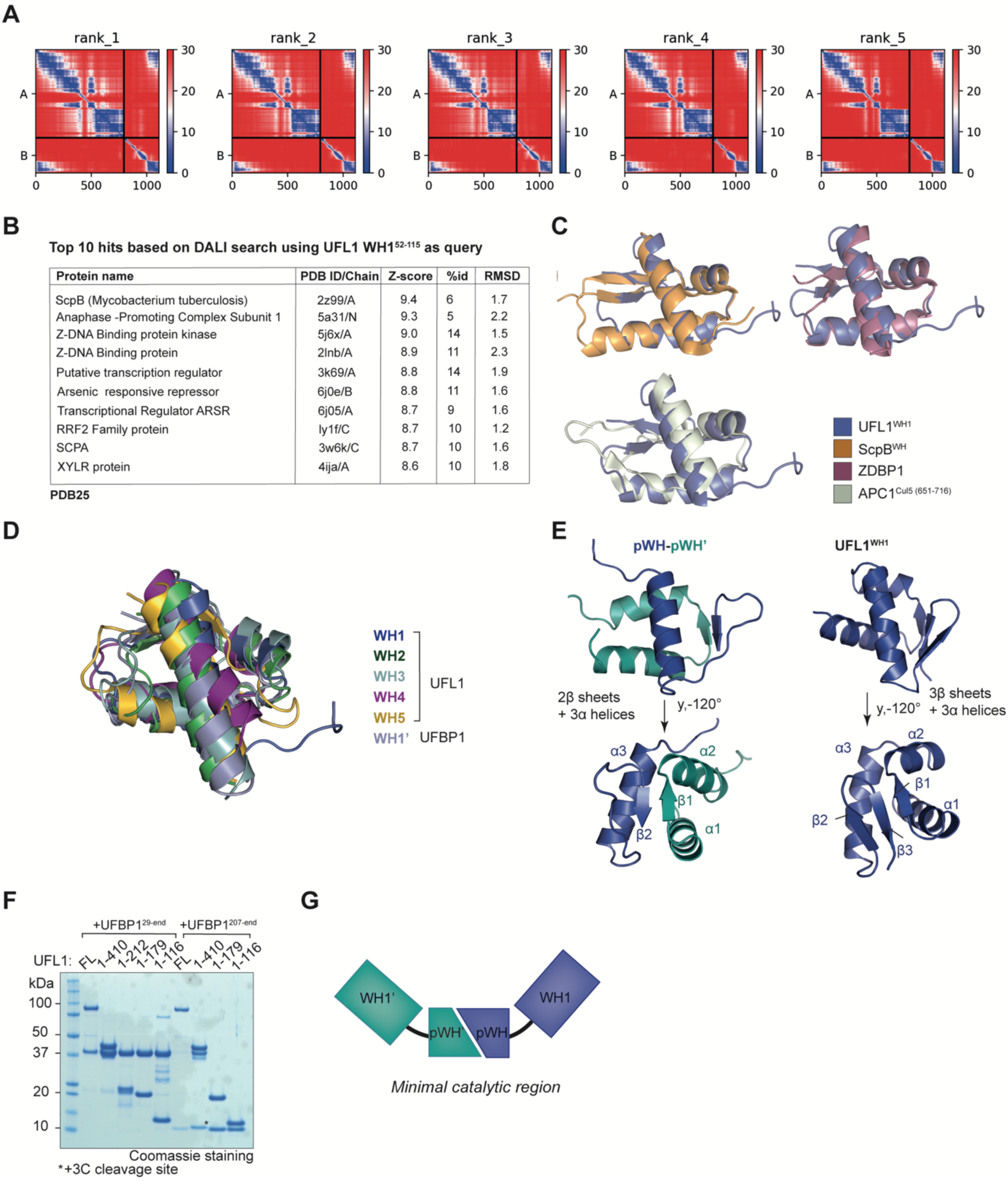
Identifying the minimal ligase domain of UFL1/UFBP1 complex. **A.** Predicted Aligned Error (PAE) scores of predicted UFL1/UFBP1 models using AlphaFold. **B.** List of structurally similar proteins from the PDB25 database predicted using UFL1 WH1(52-115) domain as query structure using DALI server. **C.** Overlaid structures of Winged helix 1(WH1) domain of UFL1(52-115) and top three hits obtained from DALI search shown in cartoon representation. **D.** Superposition of the winged helix (WH) domains of UFL1 and UFBP1. **E.** Comparison of composite winged helix domain formed by two hemi WH domains of UFL1 (pWH) and UFBP1 (pWH’) with the WH1 domain of UFL1. **F.** Coomassie stained gel showing purity of different UFL1/UFBP1 truncations. **G.** Schematic showing the tandem WH domains which constitute the minimal ligase domain.

**Figure S5:**
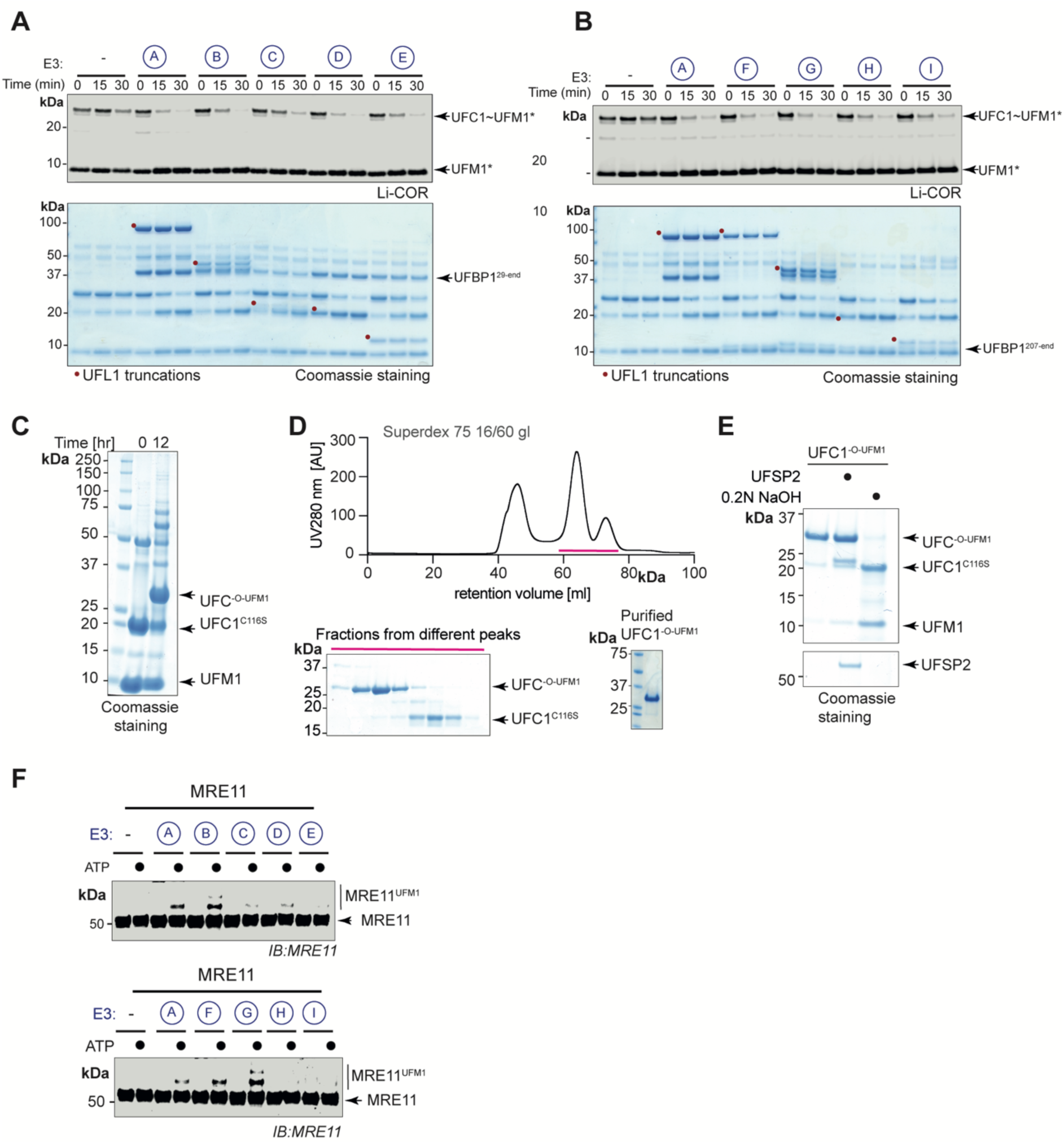
UFL1/UFBP1 E3 ligase complex binds to UFC1∼UFM1 and induces aminolysis. **A.** Lysine discharge assays to check for activation of UFC1 in the presence of different UFL1 truncations. Top gel: LiCOR scan of fluorescently labelled UFM1(UFM1*); bottom gel – Coomassie stained (Representative of three independent experiments). **B.** Lysine discharge assays as in A) in the absence of NTR of UFBP1(representative of 3 independent experiments). **C.** Coomassie stained gel showing analysis of *in vitro* UFMylation reaction products to check for the formation of stable oxy-ester linked UFC1-UFM1 conjugate (UFC1^-O-UFM1^). **D.** Chromatogram obtained from SEC analysis using HiLoad^TM^ Superdex 75 16/60pg column. (Bottom left) The fractions collected were run on an SDS-PAGE gel to identify fractions that contained pure UFC1^-O-UFM1^. (Bottom right) Coomassie stained gel showing analysis of purified UFC1^-O-UFM1^ product to check for homogeneity. **E.** Quality check to analyse if UFC1-UFM1 conjugate is linked through an oxy-ester linkage by alkaline hydrolysis. **F.** Substrate UFMylations assays to check for UFMylation of MRE11 in the presence of different UFL1/UFBP1 truncations.

**Figure S6:**
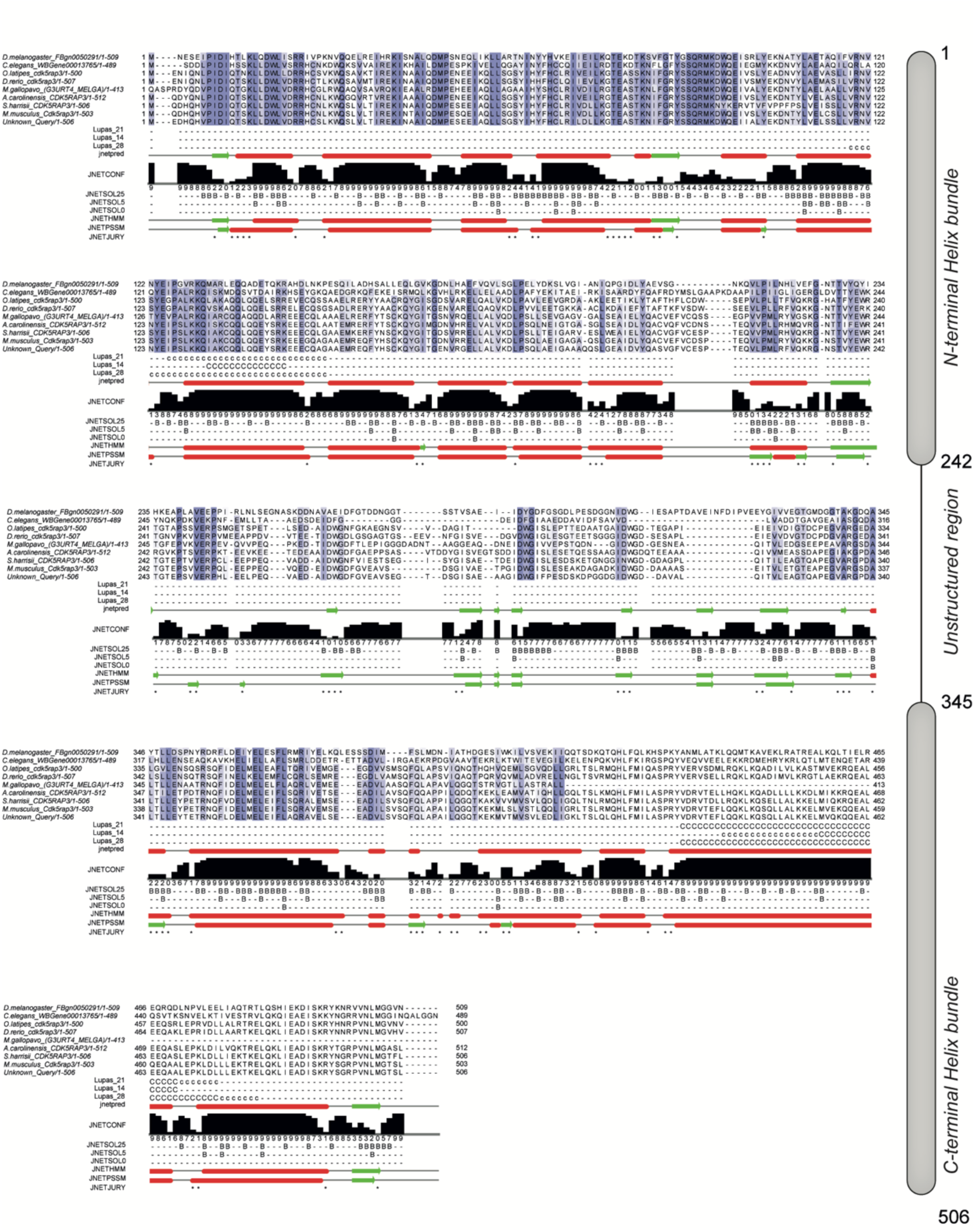
Sequence and secondary structure analysis of CDK5RAP3. **A. (Left)** Multiple sequence alignment of CDK5RAP3 homologs showing predicted secondary structure features using JPred^TM^ secondary structure program module from Jalview 2.11.1.7.**(Right)** Schematic showing predicted secondary structure features of CDK5RAP3.

**Figure S7:**
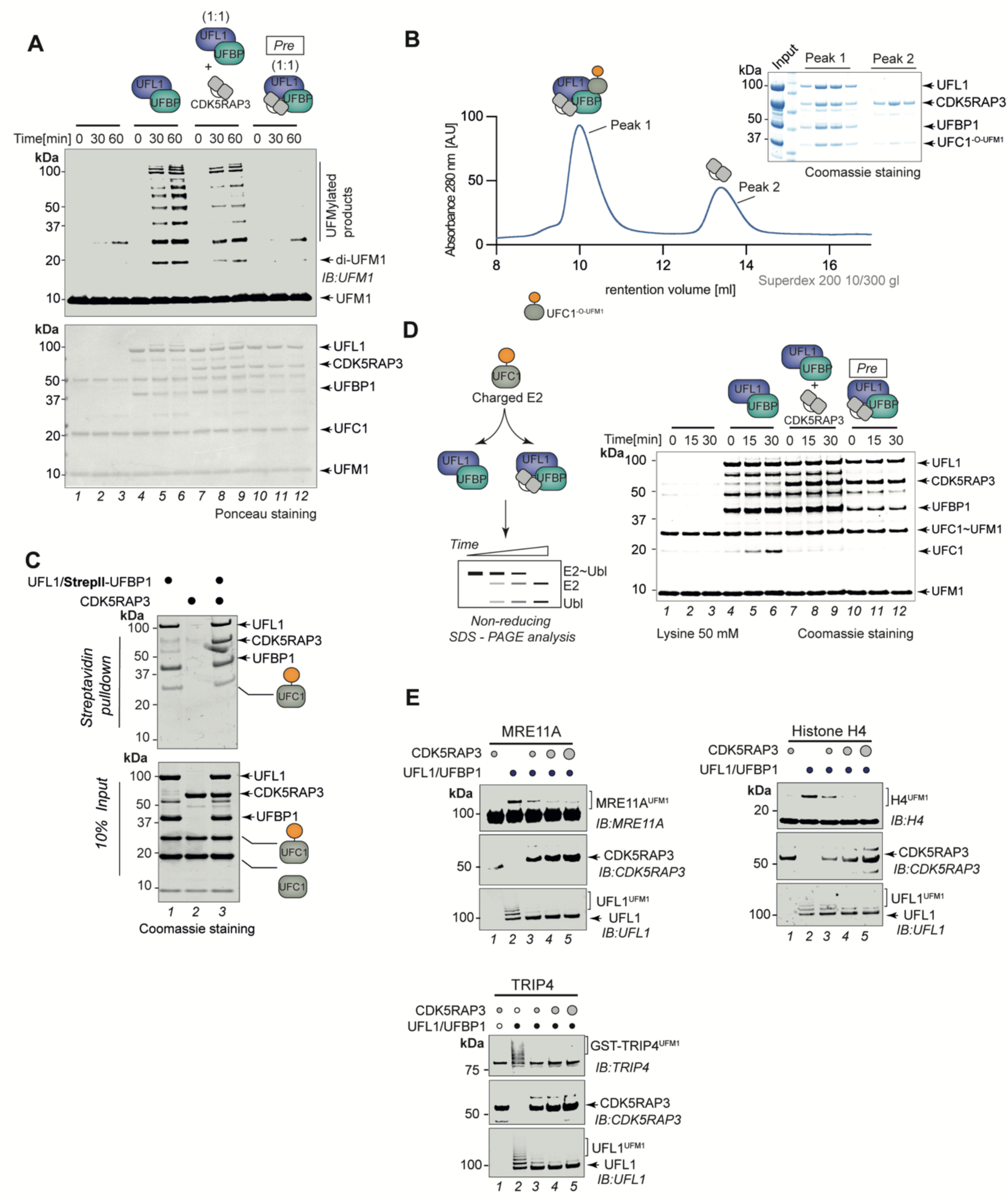
Role of CDK5RAP3 in UFMylation. **A.** *In vitro* UFMylation assay to compare the E3 ligase activity of UFL1/UFBP1 mixed with CDK5RAP3 and preassembled ternary E3 ligase complex containing UFL1/UFBP1/CDK5RAP3. Ponceau-stained membrane is showed below to indicate the amounts of reaction components used in the assay. **B.** UV traces of gel filtration chromatogram showing co-migration of (UFL1/UFBP1)/CDK5RAP3/UFC1^-O-UFM1^ complex. Approximately, 200 *µ*l of sample containing (UFL1/UFBP1)/CDK5RAP3/UFC1^-O-UFM1^ in the molar ratio of 1:1.5:3 was mixed and incubated at 4°C for 1 h and loaded onto a Superdex 200 10/300 gl column. The fractions were collected and analysed on a 4-12% SDS PAGE and visualized by Coomassie staining. **C.** Pulldown assay to check for interaction of UFL1/UFBP1 with charged UFC1 in the presence of absence of CDK5RAP3. Around 10 *µ*M of Untagged UFC1 and 10 *µ*M OF UFC1-O-UFM1 were mixed with 5 *µ*M of UFL1/UFBP1 complex in the presence and absence of CDK5RAP3. **D.** Lysine discharge assays to check for UFC1 discharge in the presence of UFL1/UFBP1 mixed with CDK5RAP3 and preassembled UFL1/UFBP1/CDK5RAP3 complex. The reaction products were run on a 4-12% SDS PAGE analysis and visualized by Coomassie staining. **E.** *In vitro* UFMylation assay to monitor UFMylation of purified substrates namely TRIP4, MRE11A and Histone H4.

**Figure S8:**
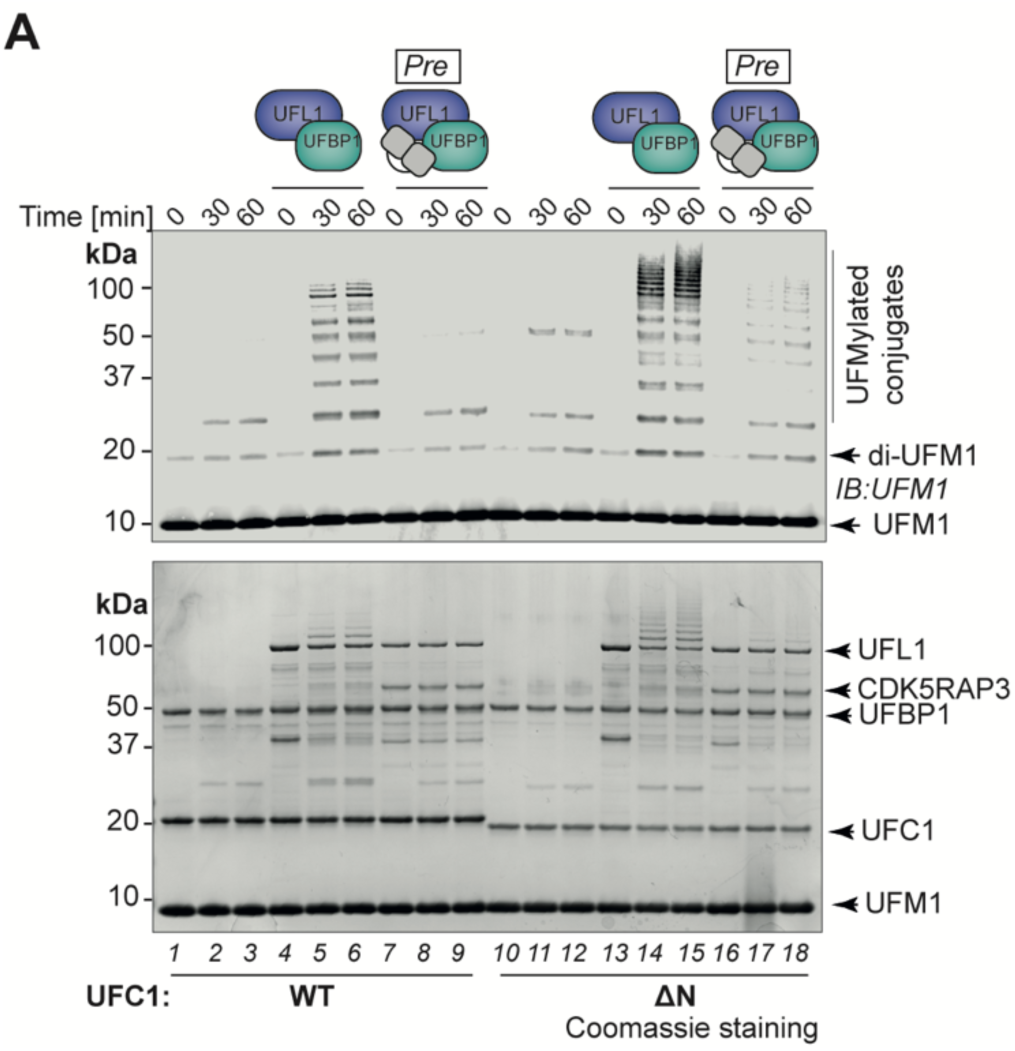
Role of the N-terminus of UFC1 in regulating UFMylation. **A.** *In vitro* UFMylation assay to compare the activites of UFC1^WT^ and UFC1^ΔN^ on its own, in the presence of UFL1/UFBP1 or in the presence of preassembled UFL1/UFBP1/CDK5RAP3 complex (Representative of three independent experiments).

**Table S1.**
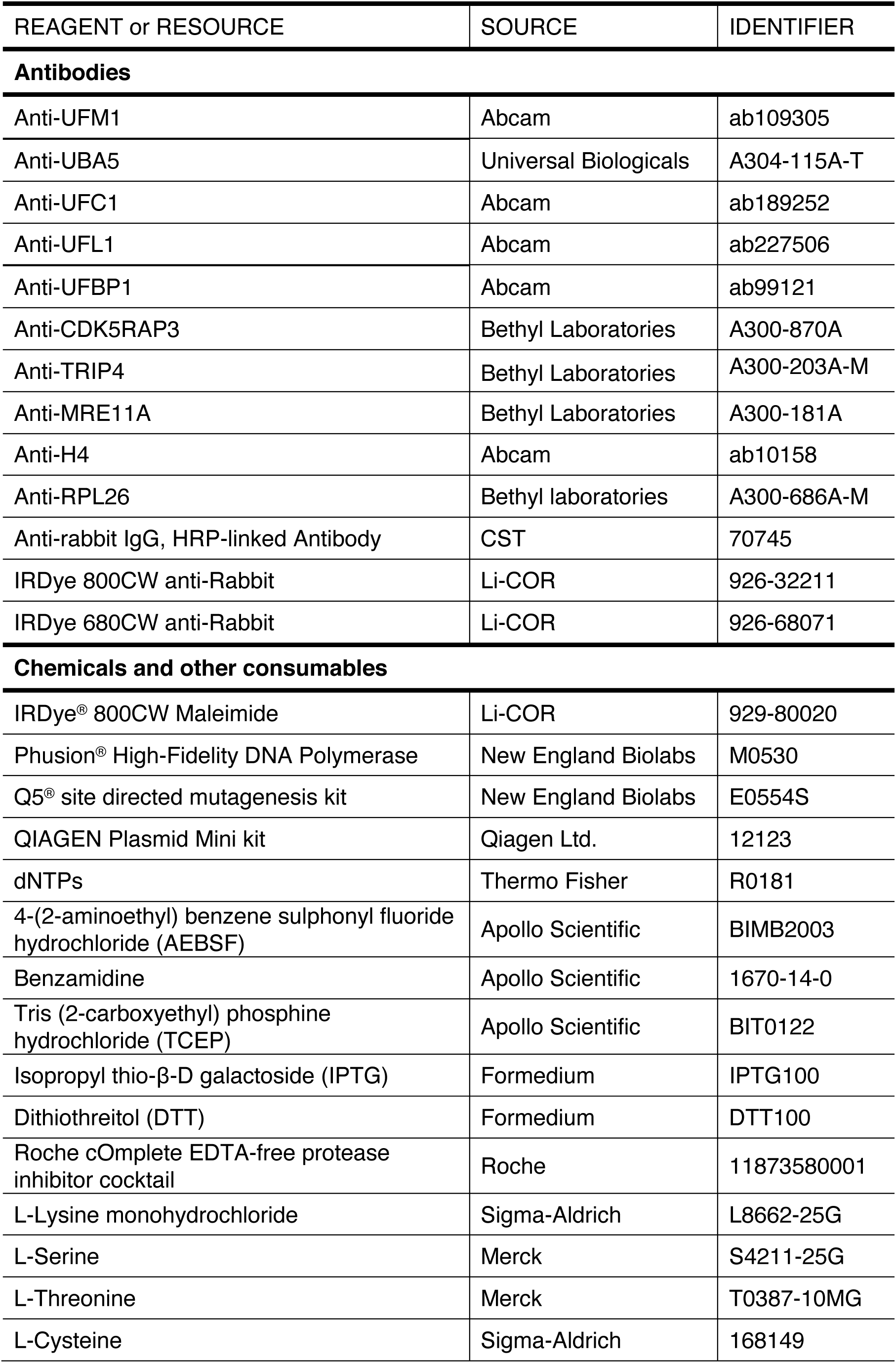
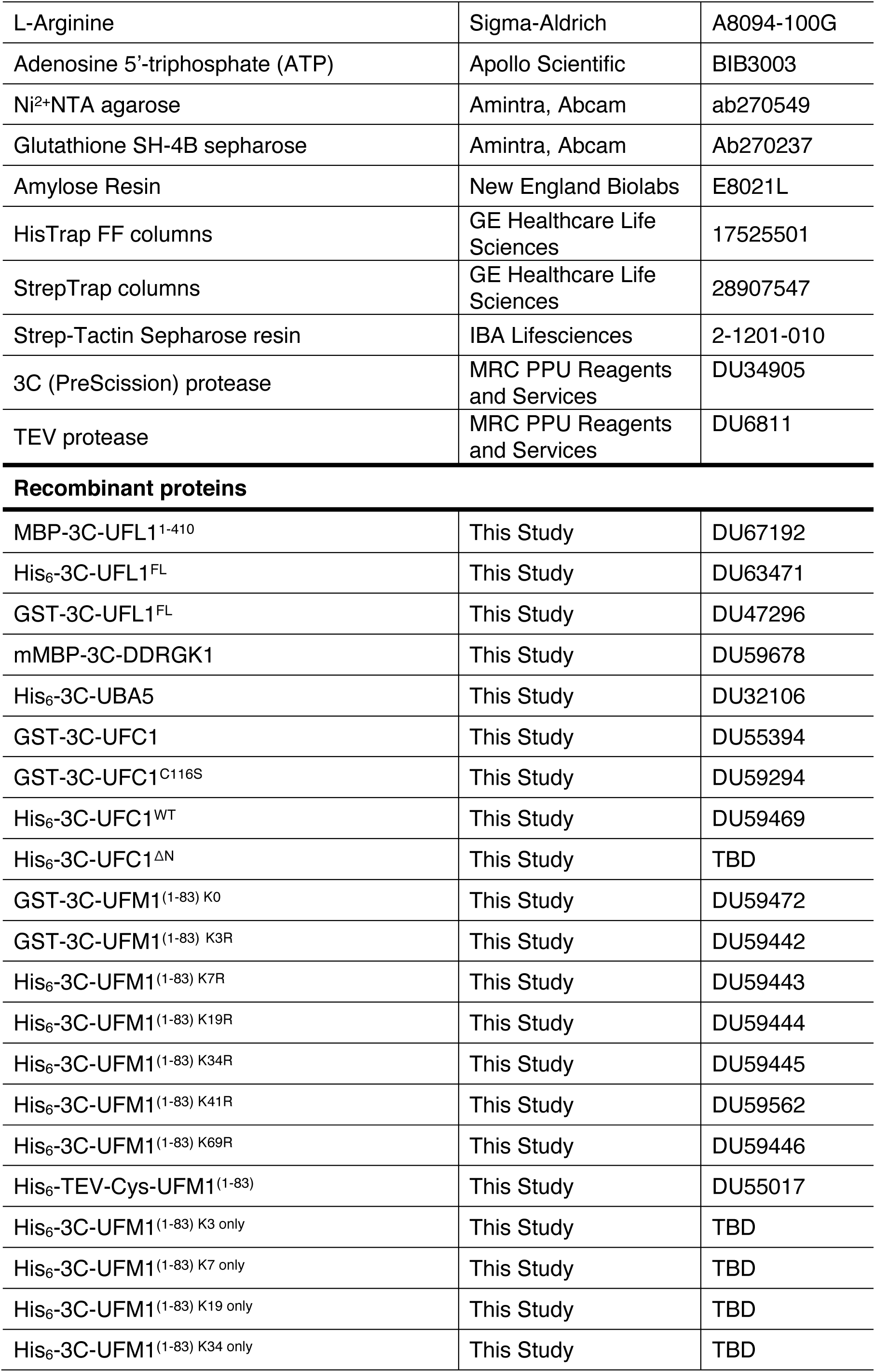
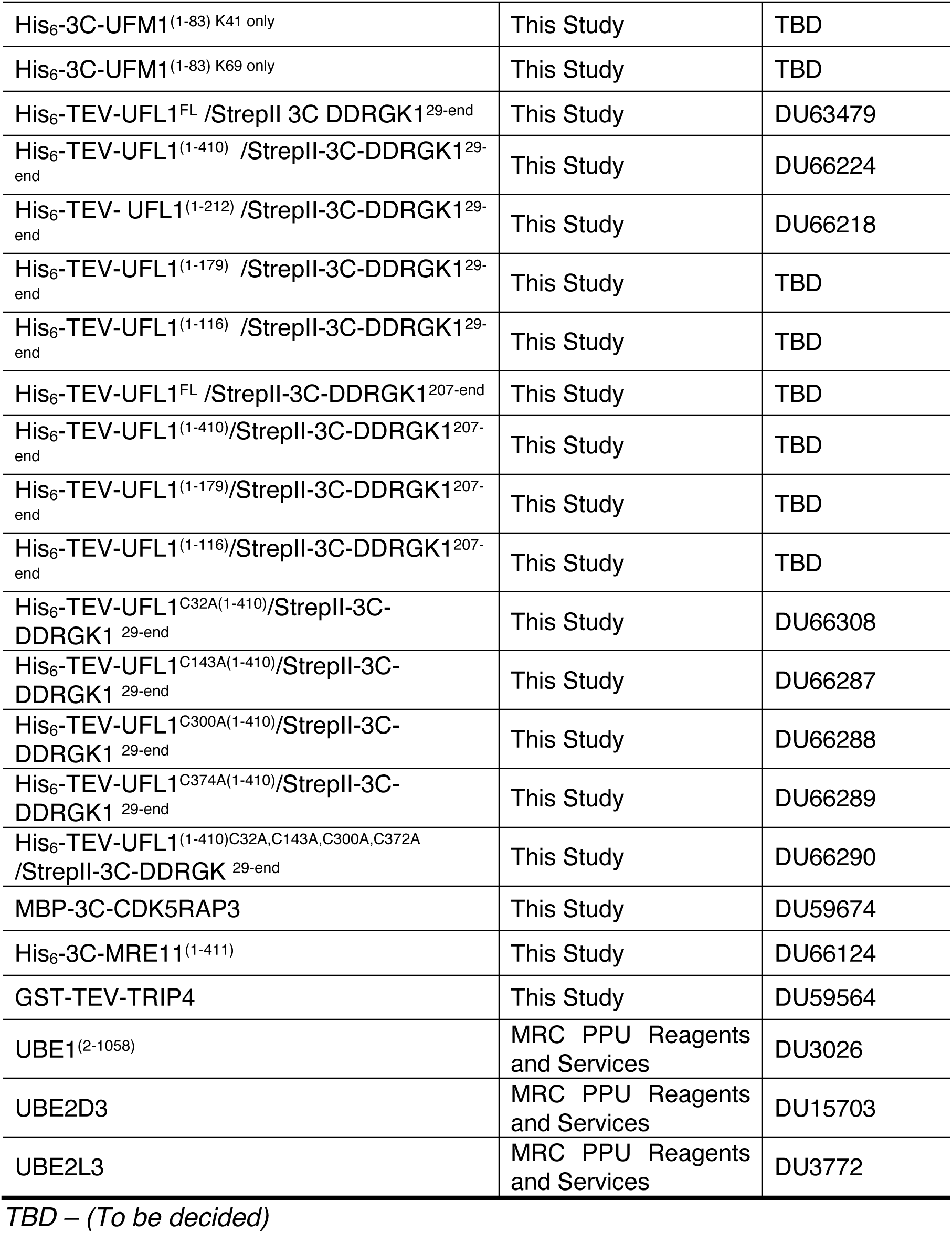

| REAGENT or RESOURCE | SOURCE | IDENTIFIER |
| --- | --- | --- |
| **Antibodies** |  |  |
| Anti-UFM1 | Abcam | ab109305 |
| Anti-UBA5 | Universal Biologicals | A304-115A-T |
| Anti-UFC1 | Abcam | ab189252 |
| Anti-UFL1 | Abcam | ab227506 |
| Anti-UFBP1 | Abcam | ab99121 |
| Anti-CDK5RAP3 | Bethyl Laboratories | A300-870A |
| Anti-TRIP4 | Bethyl Laboratories | A300-203A-M |
| Anti-MRE11A | Bethyl Laboratories | A300-181A |
| Anti-H4 | Abcam | ab10158 |
| Anti-RPL26 | Bethyl laboratories | A300-686A-M |
| Anti-rabbit IgG, HRP-linked Antibody | CST | 70745 |
| IRDye 800CW anti-Rabbit | Li-COR | 926-32211 |
| IRDye 680CW anti-Rabbit | Li-COR | 926-68071 |
| **Chemicals and other consumables** |  |  |
| IRDye<sup>®</sup> 800CW Maleimide | Li-COR | 929-80020 |
| Phusion<sup>®</sup> High-Fidelity DNA Polymerase | New England Biolabs | M0530 |
| Q5<sup>®</sup> site directed mutagenesis kit | New England Biolabs | E0554S |
| QIAGEN Plasmid Mini kit | Qiagen Ltd. | 12123 |
| dNTPs | Thermo Fisher | R0181 |
| 4-(2-aminoethyl) benzene sulphonyl fluoride hydrochloride (AEBSF) | Apollo Scientific | BIMB2003 |
| Benzamidine | Apollo Scientific | 1670-14-0 |
| Tris (2-carboxyethyl) phosphine hydrochloride (TCEP) | Apollo Scientific | BIT0122 |
| Isopropyl thio-β-D galactoside (IPTG) | Formedium | IPTG100 |
| Dithiothreitol (DTT) | Formedium | DTT100 |
| Roche cOmplete EDTA-free protease inhibitor cocktail | Roche | 11873580001 |
| L-Lysine monohydrochloride | Sigma-Aldrich | L8662-25G |
| L-Serine | Merck | S4211-25G |
| L-Threonine | Merck | T0387-10MG |
| L-Cysteine | Sigma-Aldrich | 168149 |
| L-Arginine | Sigma-Aldrich | A8094-100G |
| Adenosine 5'-triphosphate (ATP) | Apollo Scientific | BIB3003 |
| Ni<sup>2+</sup>-NTA agarose | Amintra, Abcam | ab270549 |
| Glutathione SH-4B sepharose | Amintra, Abcam | Ab270237 |
| Amylose Resin | New England Biolabs | E8021L |
| HisTrap FF columns | GE Healthcare Life Sciences | 17525501 |
| StrepTrap columns | GE Healthcare Life Sciences | 28907547 |
| Strep-Tactin Sepharose resin | IBA Lifesciences | 2-1201-010 |
| 3C (PreScission) protease | MRC PPU Reagents and Services | DU34905 |
| TEV protease | MRC PPU Reagents and Services | DU6811 |
| Recombinant proteins |  |  |
| MBP-3C-UFL1<sup>1-410</sup> | This Study | DU67192 |
| His<sub>6</sub>-3C-UFL1<sup>FL</sup> | This Study | DU63471 |
| GST-3C-UFL1<sup>FL</sup> | This Study | DU47296 |
| mMBP-3C-DDRGK1 | This Study | DU59678 |
| His<sub>6</sub>-3C-UBA5 | This Study | DU32106 |
| GST-3C-UFC1 | This Study | DU55394 |
| GST-3C-UFC1<sup>C116S</sup> | This Study | DU59294 |
| His<sub>6</sub>-3C-UFC1<sup>WT</sup> | This Study | DU59469 |
| His<sub>6</sub>-3C-UFC1<sup>ΔN</sup> | This Study | TBD |
| GST-3C-UFM1<sup>(1-83) K0</sup> | This Study | DU59472 |
| GST-3C-UFM1<sup>(1-83) K3R</sup> | This Study | DU59442 |
| His<sub>6</sub>-3C-UFM1<sup>(1-83) K7R</sup> | This Study | DU59443 |
| His<sub>6</sub>-3C-UFM1<sup>(1-83) K19R</sup> | This Study | DU59444 |
| His<sub>6</sub>-3C-UFM1<sup>(1-83) K34R</sup> | This Study | DU59445 |
| His<sub>6</sub>-3C-UFM1<sup>(1-83) K41R</sup> | This Study | DU59562 |
| His<sub>6</sub>-3C-UFM1<sup>(1-83) K69R</sup> | This Study | DU59446 |
| His<sub>6</sub>-TEV-Cys-UFM1<sup>(1-83)</sup> | This Study | DU55017 |
| His<sub>6</sub>-3C-UFM1<sup>(1-83) K3 only</sup> | This Study | TBD |
| His<sub>6</sub>-3C-UFM1<sup>(1-83) K7 only</sup> | This Study | TBD |
| His<sub>6</sub>-3C-UFM1<sup>(1-83) K19 only</sup> | This Study | TBD |
| His<sub>6</sub>-3C-UFM1<sup>(1-83) K34 only</sup> | This Study | TBD |
| His<sub>6</sub>-3C-UFM1<sup>(1-83)</sup> K41 only | This Study | TBD |
| His<sub>6</sub>-3C-UFM1<sup>(1-83)</sup> K69 only | This Study | TBD |
| His<sub>6</sub>-TEV-UFL1<sup>FL</sup> /StreptII 3C DDRGK1<sup>29-end</sup> | This Study | DU63479 |
| His<sub>6</sub>-TEV-UFL1<sup>(1-410)</sup> /StreptII-3C-DDRGK1<sup>29-end</sup> | This Study | DU66224 |
| His<sub>6</sub>-TEV- UFL1<sup>(1-212)</sup> /StreptII-3C-DDRGK1<sup>29-end</sup> | This Study | DU66218 |
| His<sub>6</sub>-TEV-UFL1<sup>(1-179)</sup> /StreptII-3C-DDRGK1<sup>29-end</sup> | This Study | TBD |
| His<sub>6</sub>-TEV-UFL1<sup>(1-116)</sup> /StreptII-3C-DDRGK1<sup>29-end</sup> | This Study | TBD |
| His<sub>6</sub>-TEV-UFL1<sup>FL</sup> /StreptII-3C-DDRGK1<sup>207-end</sup> | This Study | TBD |
| His<sub>6</sub>-TEV-UFL1<sup>(1-410)</sup>/StreptII-3C-DDRGK1<sup>207-end</sup> | This Study | TBD |
| His<sub>6</sub>-TEV-UFL1<sup>(1-179)</sup>/StreptII-3C-DDRGK1<sup>207-end</sup> | This Study | TBD |
| His<sub>6</sub>-TEV-UFL1<sup>(1-116)</sup>/StreptII-3C-DDRGK1<sup>207-end</sup> | This Study | TBD |
| His<sub>6</sub>-TEV-UFL1<sup>C32A(1-410)</sup>/StreptII-3C-DDRGK1<sup>29-end</sup> | This Study | DU66308 |
| His<sub>6</sub>-TEV-UFL1<sup>C143A(1-410)</sup>/StreptII-3C-DDRGK1<sup>29-end</sup> | This Study | DU66287 |
| His<sub>6</sub>-TEV-UFL1<sup>C300A(1-410)</sup>/StreptII-3C-DDRGK1<sup>29-end</sup> | This Study | DU66288 |
| His<sub>6</sub>-TEV-UFL1<sup>C374A(1-410)</sup>/StreptII-3C-DDRGK1<sup>29-end</sup> | This Study | DU66289 |
| His<sub>6</sub>-TEV-UFL1<sup>(1-410)C32A,C143A,C300A,C372A</sup> /StreptII-3C-DDRGK<sup>29-end</sup> | This Study | DU66290 |
| MBP-3C-CDK5RAP3 | This Study | DU59674 |
| His<sub>6</sub>-3C-MRE11<sup>(1-411)</sup> | This Study | DU66124 |
| GST-TEV-TRIP4 | This Study | DU59564 |
| UBE1<sup>(2-1058)</sup> | MRC PPU Reagents and Services | DU3026 |
| UBE2D3 | MRC PPU Reagents and Services | DU15703 |
| UBE2L3 | MRC PPU Reagents and Services | DU3772 |

TBD – (To be decided)

## Notes

### Competing Interest Statement

The authors have declared no competing interest.

